# Comparative Neuroanatomy of the Lumbosacral Spinal Cord of the Rat, Cat, Pig, Monkey, and Human

**DOI:** 10.1101/2020.08.11.246165

**Authors:** Amirali Toossi, Bradley Bergin, Maedeh Marefatallah, Behdad Parhizi, Neil Tyreman, Dirk G. Everaert, Sabereh Rezaei, Peter Seres, J. Christopher Gatenby, Steve I. Perlmutter, Vivian K. Mushahwar

**Affiliations:** Krembil Research Institute, University Health Network, Toronto, Canada; Department of Medicine, University of Alberta, Edmonton, Canada; Department of Chemical and Materials Engineering, University of Alberta, Edmonton, Canada; Neuroscience and Mental Health Institute, University of Alberta, Edmonton, Canada; Department of Materials Science and Engineering, University of Toronto, Toronto, Canada; Department of Biomedical Engineering, University of Alberta, Edmonton, Canada; Department of Radiology, University of Washington, Seattle, USA; Department of Physiology and Biophysics, University of Washington, Seattle, USA; Washington National Primate Research Centre, Seattle, USA; Sensory Motor Adaptive Rehabilitation Technology (SMART) Network, University of Alberta, Edmonton, Canada

**Keywords:** Spinal cord, Lumbosacral Enlargement, Human, Monkey, Swine, Feline, Rodent, Atlas, Anatomy, Morphology, Magnetic resonance imaging, Intraspinal delivery, Intraparenchymal delivery, spinal cord disease, spinal cord injury

## Abstract

The overall goal of this work was to create a high-resolution MRI atlas of the lumbosacral enlargement of the spinal cord of the rat (Sprague-Dawley), cat, domestic pig, rhesus monkey, and human. These species were chosen because they are commonly used in basic and translational research in spinal cord injuries and diseases. Six spinal cord specimens from each of the studied species (total of 30 specimens) were fixed, extracted, and imaged. Sizes of the spinal cord segments, cross-sectional dimensions, and locations of the spinal cord gray and white matter were quantified and compared across species. The obtained atlas establishes a reference for the neuroanatomy of the intact lumbosacral spinal cord in these species. It can also be used to guide the planning of surgical procedures of the spinal cord, technology design and development of spinal cord neuroprostheses, and the precise delivery of cells/drugs into target regions within the spinal cord parenchyma.

## Introduction

The spinal cord is an important part of the central nervous system that plays an essential role in the function of the body’s sensorimotor and autonomic systems. Spinal cord injuries and diseases (e.g., spinal cord injury [1], spinal muscular atrophy [2], and amyotrophic lateral sclerosis [3]) affect the lives of millions of people around the world, impacting their physical and mental health as well as the socioeconomical aspects of their lives [4]–[9]. Detailed knowledge of the neuroanatomy of the spinal cord is critical for understanding its pathologies and finding possible cures. Because many investigations into spinal cord injuries and diseases involve animal models, appropriate selection of an animal model and correct interpretation of study findings requires an understanding of the anatomical and physiological similarities and differences of the models in relation to humans.

The neuroanatomy of the spinal cord has been a subject of curiosity for centuries, dating back to the third and fourth centuries B.C.E., and including observations made by Hippocrates [10]. Since then, numerous studies have been conducted on the anatomy and physiology of the spinal cord in various species. However, anatomical atlases of the spinal cord are still limited in number and commonly have a coarse spatial resolution and/or are limited to a single species. One of the first spinal cord atlases was published by Alexander Bruce, in 1901. This Atlas provided transverse sections of the entire human spinal cord with 1 section per spinal cord segment [11]. In 2008, the *Allen Institute* published a histological spinal cord atlas for the mouse [12] along with results from Nissl staining and RNA in situ hybridization. This atlas has a longitudinal resolution of 2 mm and spans the entire spinal cord of 4- and 56-days postnatal animals. Tokuno et al created a spinal cord atlas for the macaque monkey, with a resolution of 1 transverse section per spinal cord segment [13]. Most recently, Sengul et al created a multispecies spinal cord atlas for the rat, mouse, marmoset, rhesus monkey and human. This atlas spans the entire length of the spinal cord for each species, but also has a resolution of only 1 transverse section per spinal cord segment [14].

Recent advances in the development of targeted therapies that require precise access to regions within the central nervous system highlight the importance of high-resolution anatomical atlases. Examples include deep brain stimulation implants for Parkinson’s disease [15], epidural [16] and intraspinal microstimulation for restoring movement after severe spinal cord injury [17]–[19], and stem cell transplantation treatments for amyotrophic lateral sclerosis [20]. These interventions commonly employ stereotactic techniques guided by electrophysiology, imaging and knowledge from anatomical atlases to further enhance their anatomical targeting and treatment effectiveness [21]–[23]. To address the need for increased anatomical detail of the spinal cord for intraspinal therapeutics, this study aimed to complement existing knowledge of spinal cord neuroanatomy by creating a comparative high-resolution spinal cord atlas for species commonly utilized in preclinical and translational research [24]. We used magnetic resonance imaging (MRI) to image and analyze the spinal cords of rats, cats, domestic pigs, rhesus monkeys, and humans, and created atlases with a minimum resolution of 1 mm x 0.15 mm x 0.15 mm in the longitudinal, sagittal and transverse axes, respectively. This work was focused on studying the region of the lumbosacral enlargement of the spinal cord which houses a dense neuronal network involved in the control of motor and sensory functions of the lower limbs (e.g., standing and walking [25], [26]), and autonomic functions (e.g., modulation of blood pressure [27]). The resulting comparative atlas provides necessary information for the development, implementation and translation of novel and precise treatments of spinal cord injuries and diseases.

## Results

MRI atlases were created based on spinal cord specimens extracted from 30 cadavers: 6 Sprague-Dawley rats, 6 cats, 6 rhesus macaque monkeys, 6 domestic pigs, and 6 humans (Table 1).

**Table 1.** Metadata for the spinal cord specimens used in this study. N/A – not applicable. * – approximate age of the domestic pigs can be estimated from available porcine growth rate charts [52], [53].

| Sample ID | Species | Strain | Sex | Age (years) | Weight (kg) | Height (m) |
| --- | --- | --- | --- | --- | --- | --- |
| R1 | Rat | Sprague Dawley | Female | 0.5 | 320 | N/A |
| R2 | Rat | Sprague Dawley | Female | 0.5 | 345 | N/A |
| R3 | Rat | Sprague Dawley | Female | 0.5 | 360 | N/A |
| R4 | Rat | Sprague Dawley | Female | 0.5 | 350 | N/A |
| R5 | Rat | Sprague Dawley | Female | 0.5 | 400 | N/A |
| R6 | Rat | Sprague Dawley | Female | 0.5 | 380 | N/A |
| C1 | Cat | N/A | Male | 7.7 | 4.7 | N/A |
| C2 | Cat | N/A | Male | 4.4 | 4.7 | N/A |
| C3 | Cat | N/A | Female | 7.4 | 3.22 | N/A |
| C4 | Cat | N/A | Female | 7.3 | 5.86 | N/A |
| C5 | Cat | N/A | Female | 1.7 | 4.24 | N/A |
| C6 | Cat | N/A | Female | 1.6 | 5.9 | N/A |
| P1 | Pig | Domestic | Female | * | 50.9 | N/A |
| P2 | Pig | Domestic | Female | * | 57.4 | N/A |
| P3 | Pig | Domestic | Female | * | 49 | N/A |
| P4 | Pig | Domestic | Female | * | 42 | N/A |
| P5 | Pig | Domestic | Male | * | 42 | N/A |
| P6 | Pig | Domestic | Female | * | 43 | N/A |
| M1 | Monkey | Rhesus Macaque | Male | Not available | 13.2 | N/A |
| M2 | Monkey | Rhesus Macaque | Female | 18.7 | 6.85 | N/A |
| M3 | Monkey | Rhesus Macaque | Female | 16.9 | 8 | N/A |
| M4 | Monkey | Rhesus Macaque | Male | 9 | 15.1 | N/A |
| M5 | Monkey | Rhesus Macaque | Female | Not available | 10.5 | N/A |
| M6 | Monkey | Rhesus Macaque | Female | 18.9 | 11.5 | N/A |
| H1 | Human | N/A | Male | 69 | Not available | 1.78 |
| H2 | Human | N/A | Female | 91 | Not available | 1.65 |
| H3 | Human | N/A | Male | 79 | Not available | 1.78 |
| H4 | Human | N/A | Male | 84 | Not available | Not available |
| H5 | Human | N/A | Male | 91 | Not available | 1.62 |
| H6 | Human | N/A | Male | 86 | Not available | 1.76 |

### Spinal Cord Segments and Longitudinal Dimensions

The total length of the L1-S1 region was largest in pigs (116.3±5 mm, p<0.001 for all comparisons), followed by cats (92.1±3.9 mm, p<0.001 for all comparisons), monkeys (65±3.9 mm, p<0.001 for all comparisons except relative to humans), humans (61.4±4.5 mm, p<0.001 for all comparisons except relative to monkeys), and rats (19.8±2.3 mm, p<0.001 for all comparisons), respectively. In all species, except in rats, spinal cord segments gradually become shorter in length, moving from the rostral to the caudal end (Fig. 1).

**Fig. 1.**
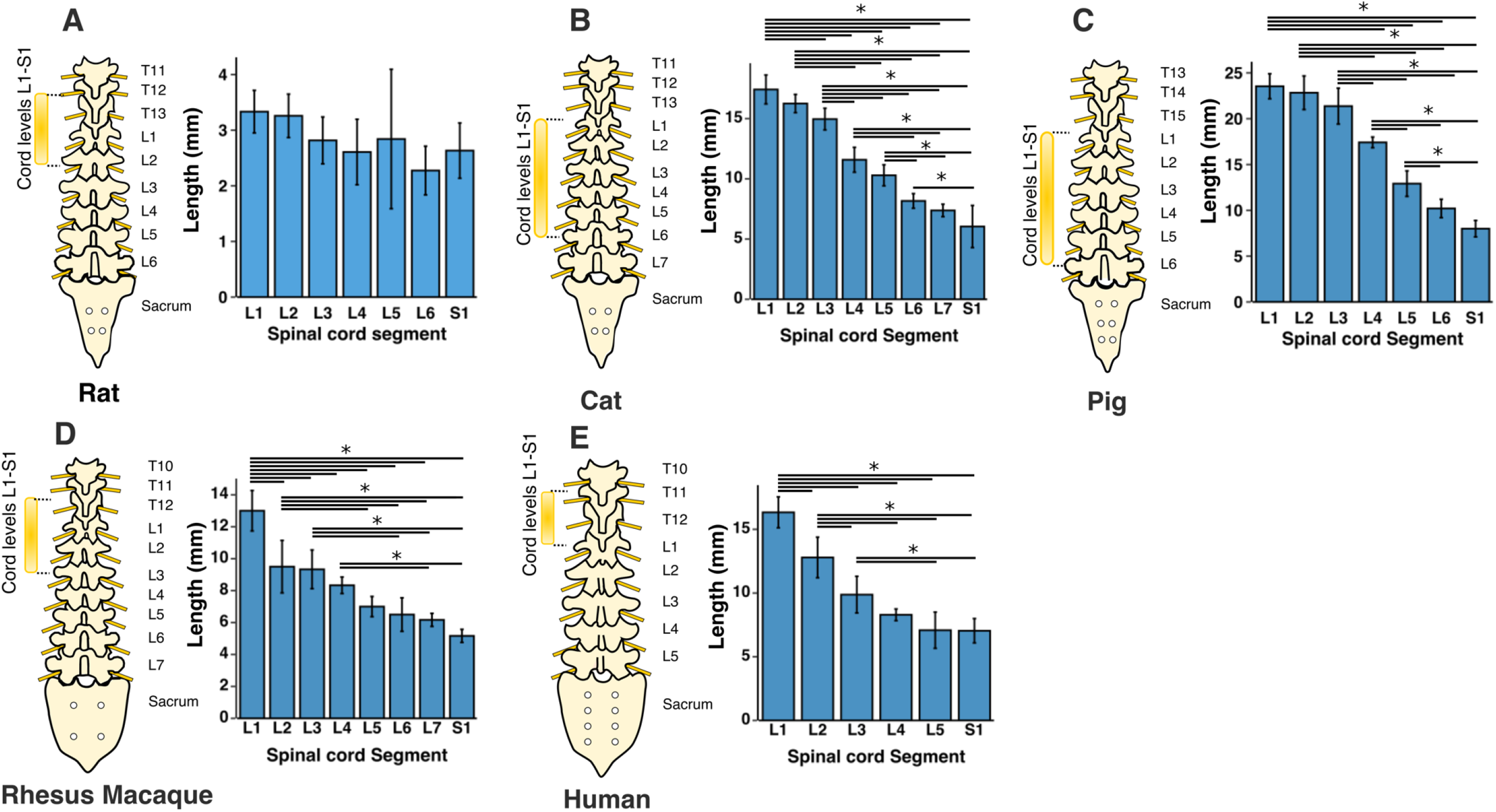
Segmental length of the spinal cord levels L1-S1. Bars represent the mean and the error bars show the standard deviation of the mean. * represents p<0.05.

The lumbosacral enlargement was defined as the region of the spinal cord that houses the motoneurons that innervate the muscles of the lower extremities. This region was identified based on the morphological features of the gray matter described by Vanderhorst and Holstege [28] and Gross et al [29], as shown in Fig. 2. Accordingly, moving caudally from the rostral end of the lumbar enlargement, the ventral horns of the gray matter become larger and gradually extrude in the lateral direction. The opposite occurs at the caudal end of the enlargement, where the ventral horns become smaller and gradually contract towards the medial direction. Using this identification method, the lumbosacral enlargement typically spanned spinal cord segments L3-S1 in rats, L4-S1 in cats, L3-S1 in pigs, L2/L3-L7/S1 in monkeys, and T12/L1-S1/S2 in humans (Fig. 3 and Table S1), consistent with findings from the literature [28], [30]–[37]. The enlargement is located in the low thoracic and high lumbar regions of the vertebral column in rats, monkeys, and humans, while in cats and pigs it is positioned in the lower lumbar region of the spine (Fig. 3A-3E). The length of the enlargement was longest in pigs (66.6±5.6 mm, p<0.05 for all comparisons), followed by humans (57.6±5.1 mm, p<0.05 for all comparisons), followed by cats and monkeys (34.3±1.5 mm and 34.8±5.7 mm, respectively, p<0.001 for all comparisons except relative to each other where p=0.99), followed by rats (12.3±2.4 mm, p<0.001 for all comparisons) (Fig. 3F).

**Fig. 2.**
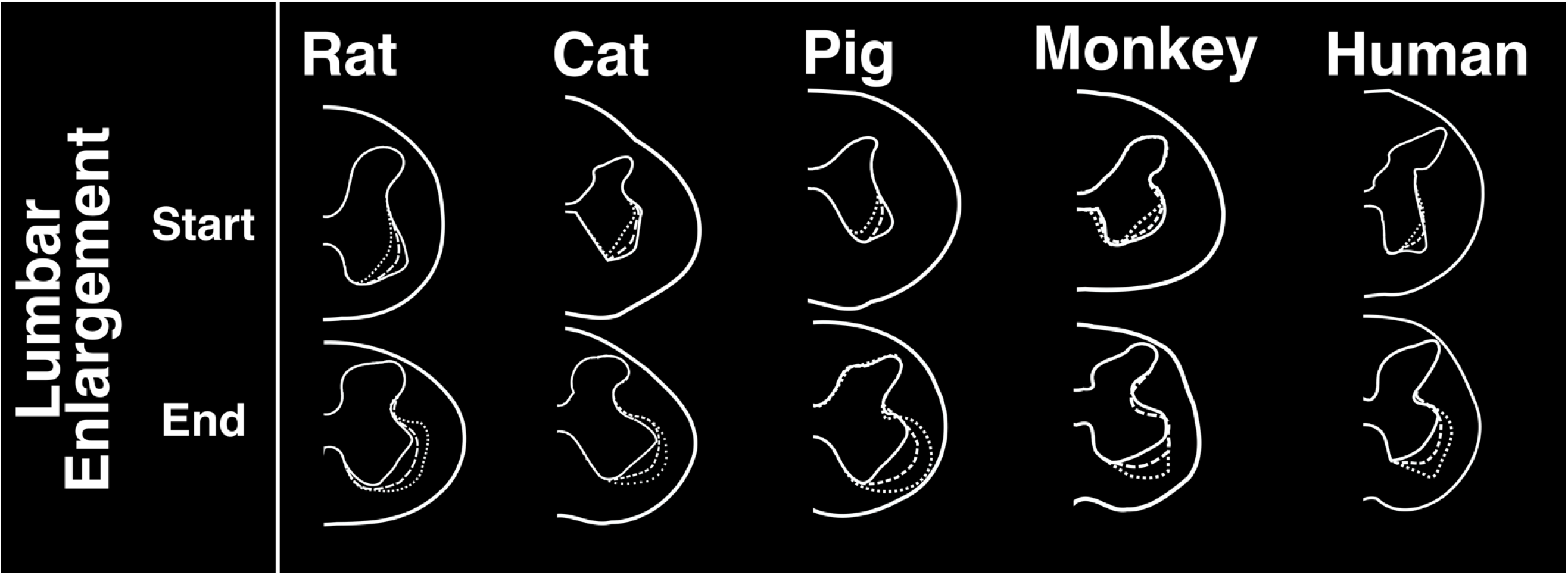
Characteristic morphological changes of the gray matter of the spinal cord at the rostral and caudal boundaries of the lumbosacral enlargement. These morphological changes were described by Vanderhorst and Holstege [28] and Gross et al [29]. Cross-sectional shapes illustrated here were derived from representative MRI images in each species. Dotted, dashed and solid traces show morphological changes of the gray matter moving from rostral to caudal direction, respectively.

**Fig. 3.**
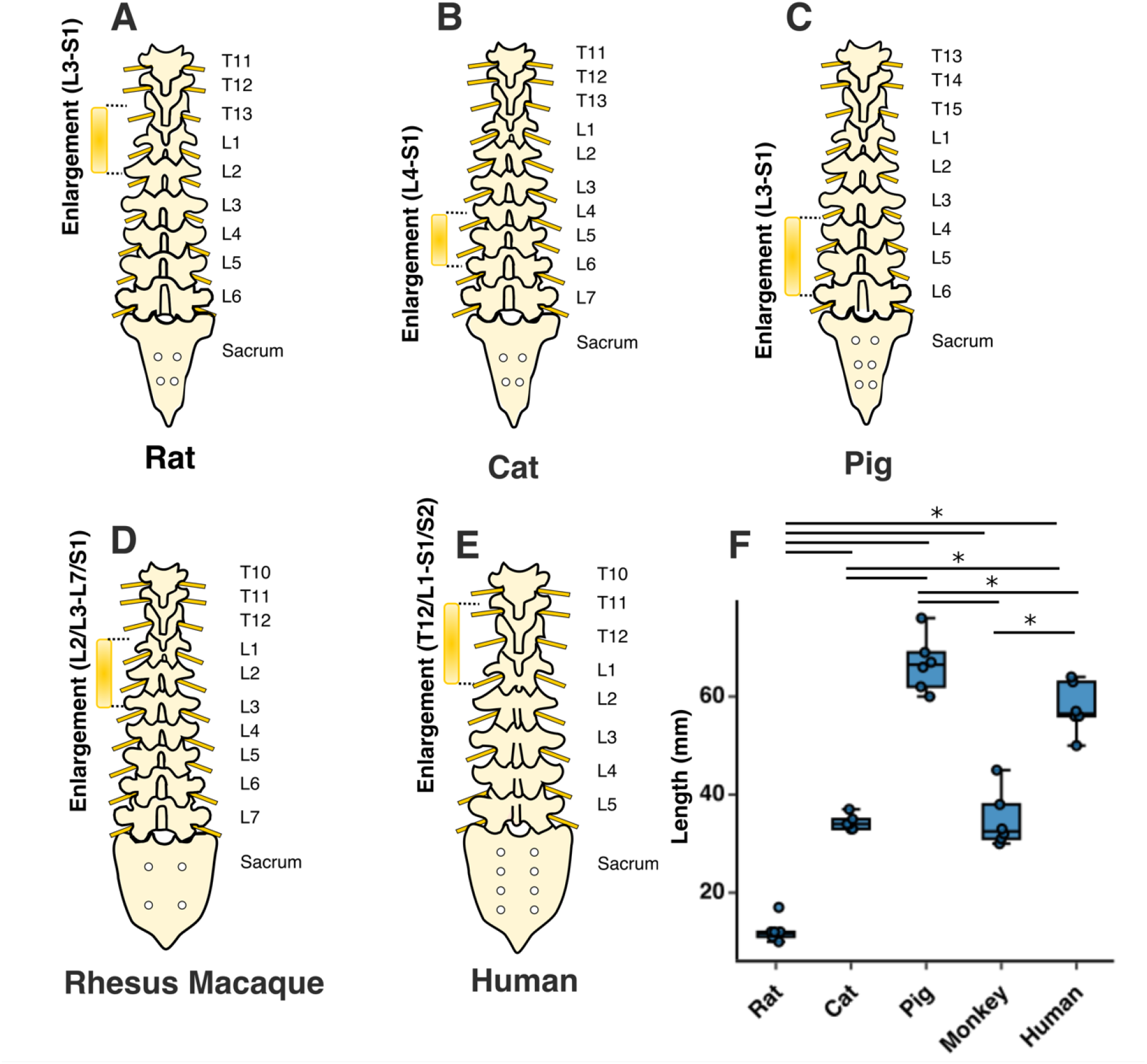
Rostrocaudal extent and location of the lumbosacral enlargement in various species. The typical segmental levels of the lumbosacral enlargement were: L3-S1 for rats, L4-S1 for cats, L3-S1 for pigs, L2/L3-L7/S1 for rhesus macaque, and T12/L1-S1/S2 for humans. Levels for enlargement (in brackets) are spinal cord segments; the levels on the right side of the spinal column indicate the vertebral levels. Boxes represent interquartile range; horizontal line shows the median; whiskers represent minimum and maximum values of the dataset. ‘*’ symbol represents p<0.05. Solid dots show individual data points.

### Gray and White Matter and Cross-sectional Dimensions

For all species, representative cross-sectional images from each spinal segment within the lumbosacral spinal cord are shown in Fig. 4, and MRI-based 3D reconstructed models are shown in Fig. 5. Serial cross-sectional images of the enlargement are shown in Fig. 6.

**Fig 4.**
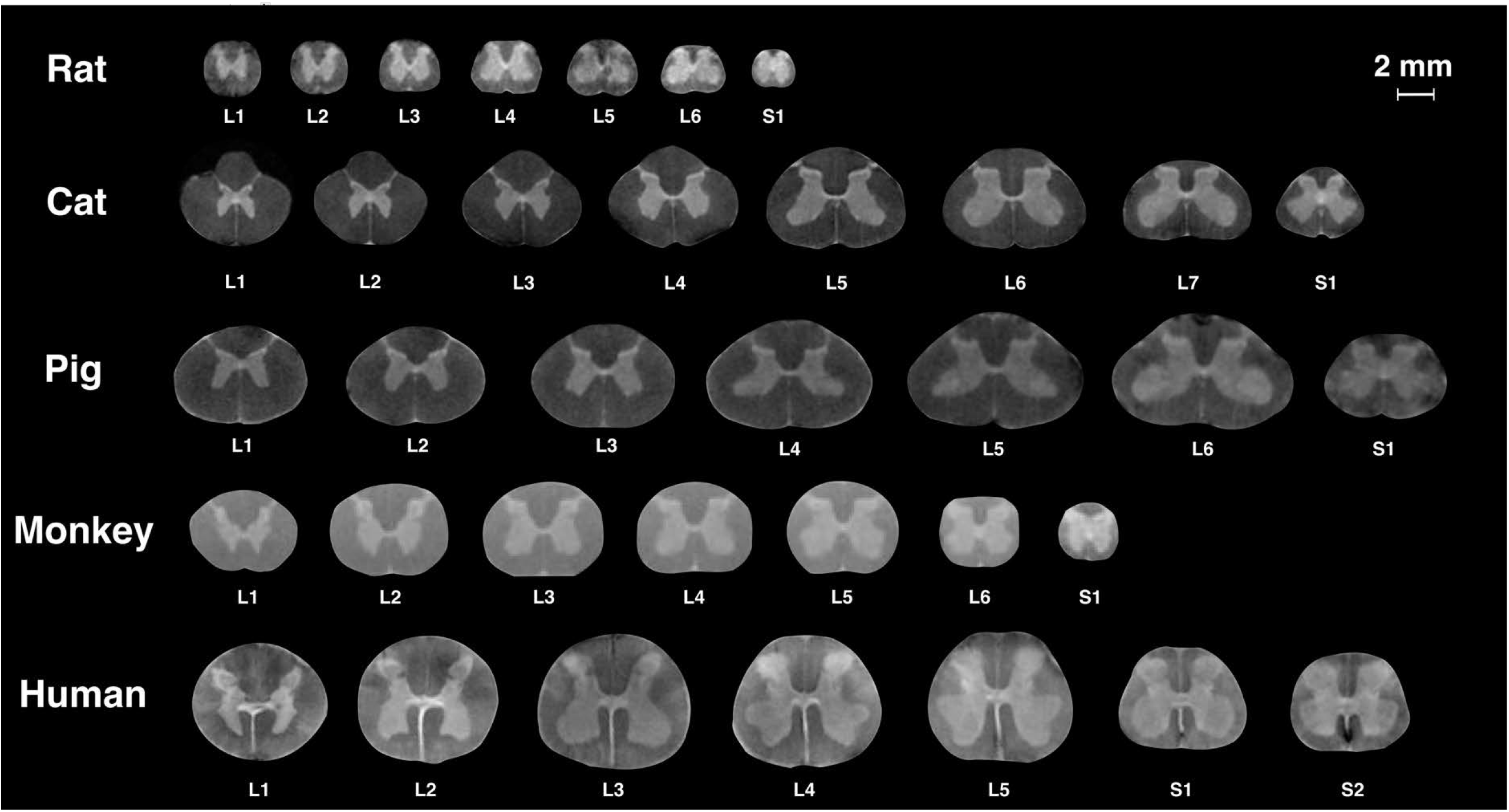
MR images of spinal cord segments L1-S1. Images of the spinal cords of rats, cats, pigs, and humans were acquired using a 4.7T scanner. Images of the spinal cords of monkeys were acquired using a 3T scanner. Each cross-sectional image is taken from the middle of the corresponding spinal cord segment.

**Fig 5.**
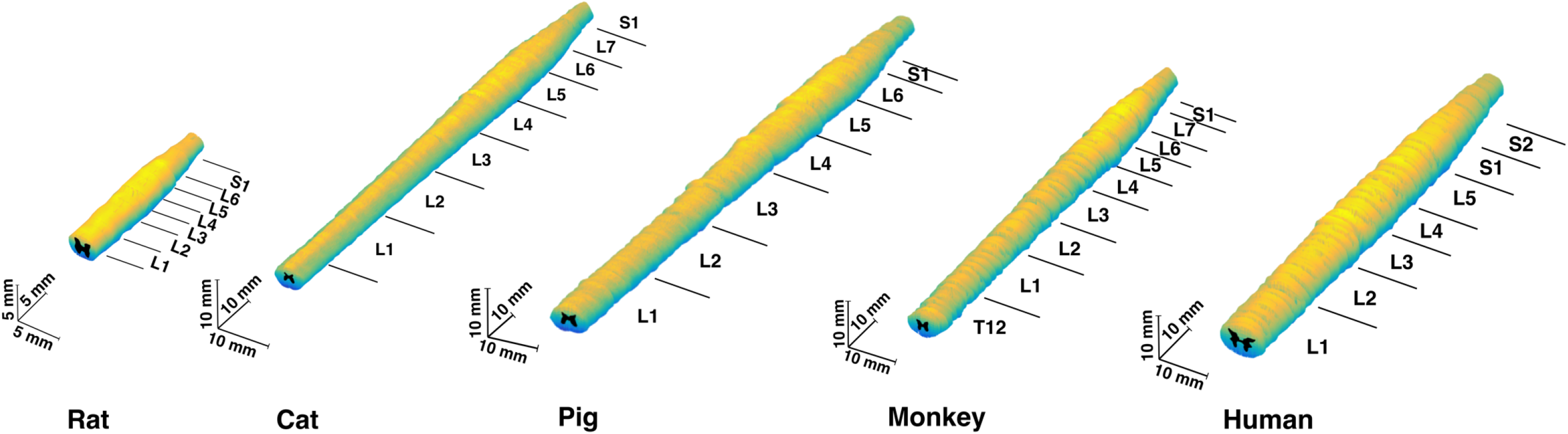
3D model of the lumbosacral spinal cords of the rat, cat, pig, monkey, and human. 3D models were reconstructed based on the acquired MRIs from a representative animal (n=1) per species. Annotations show the segments of the spinal cord identified using the method shown in Fig.S7.

**Fig 6.**
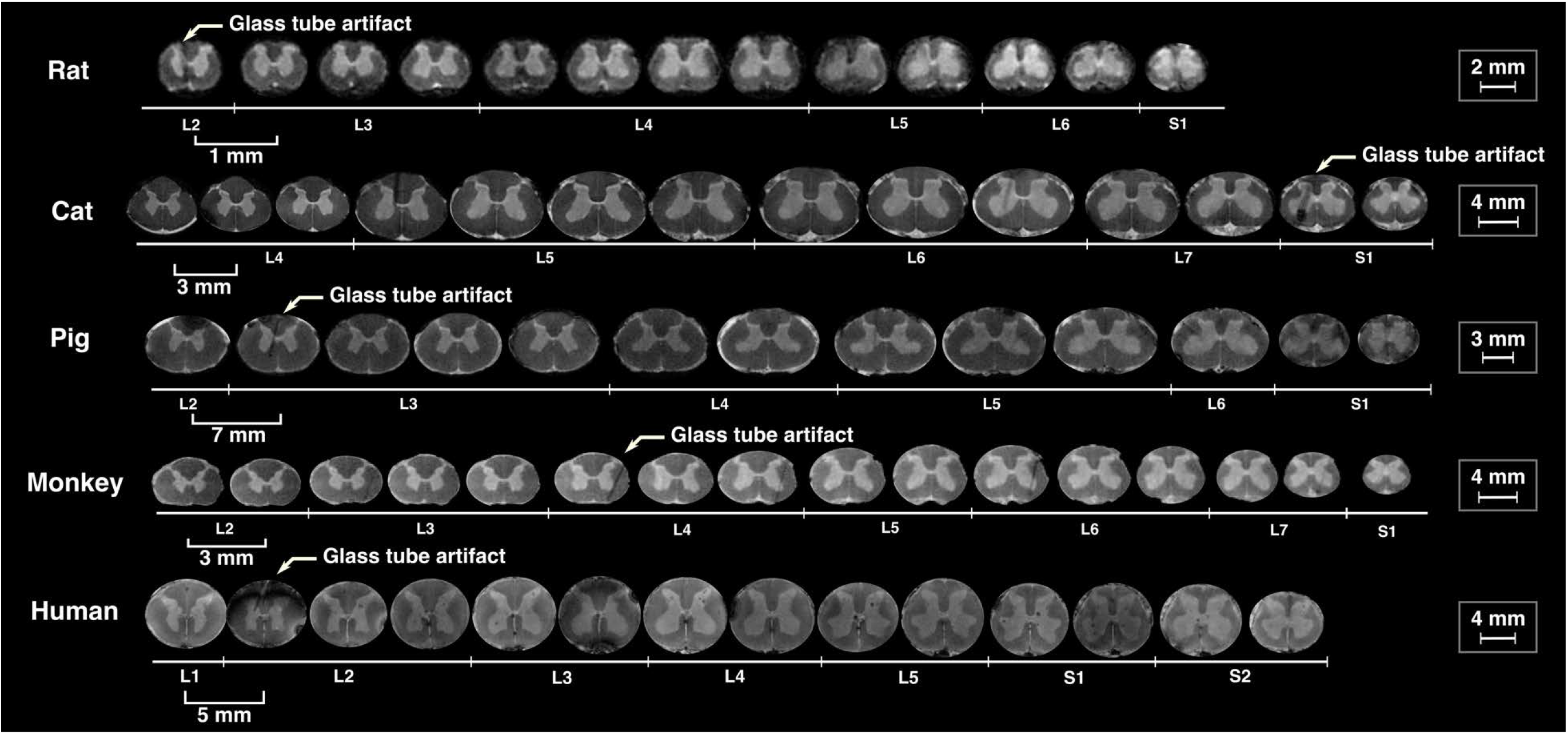
Sequential cross sections from the lumbosacral enlargement of the spinal cords of rat, cat, pig, monkey, and human, spaced apart by 1, 3, 7, 3 and 5 mm, respectively. Scale bars for each species are shown on the right. The artifacts (arrows) are the locations of the glass markers. Images of the spinal cords of rats, cats, pigs, and humans were acquired using a 4.7T scanner. Images of the spinal cords of monkeys were acquired using a 3T scanner.

The morphologies of the gray and white matter of the spinal cord were quantified (Fig. 7) to document their changes along the longitudinal (rostrocaudal) axis for all species (Fig. 8–12). Measurements for each individual specimen are shown in supplementary Figs. S1–S5. Table 2 shows the intraclass correlation (ICC) calculations for these measurements. Reliability of the measurements was good or excellent for all measured parameters except for d6 (Fig. 7, middle), which was moderate.

**Table 2.** Intraclass Correlation (ICC) calculations as a measure of inter-rater reliability of the measurements made based on MRIs. Presented ICC values are the *average score ICCs* calculated based on the one-way random model. CI: 95% confidence interval for ICC calculations.

| Parameters | Unit | Total Number of Raters | ICC(A,2), p-Value [95% CI] | Reliability Assessment |
| --- | --- | --- | --- | --- |
| d1 | mm | 3 | 0.985, p<0.001<br>[0.984, 0.986] | Excellent |
| d2 | mm | 3 | 0.973, p<0.001<br>[0.97, 0.975] | Excellent |
| d3 | mm | 3 | 0.991, p<0.001<br>[0.99, 0.992] | Excellent |
| d4 | mm | 3 | 0.977, p<0.001<br>[0.975, 0.979] | Excellent |
| d5 | mm | 3 | 0.979, p<0.001<br>[0.977, 0.98] | Excellent |
| d6 | mm | 3 | 0.734, p<0.001<br>[0.712, 0.755] | Moderate |
| d7 | mm | 3 | 0.948, p<0.001<br>[0.944, 0.952] | Excellent |
| d8 | mm | 3 | 0.831, p<0.001<br>[0.817, 0.844] | Good |
| d9 | mm | 3 | 0.911, p<0.001<br>[0.902, 0.919] | Excellent |
| a1 | mm<sup>2</sup> | 3 | 0.995, p<0.001<br>[0.994, 0.995] | Excellent |
| a2 | mm<sup>2</sup> | 2 | 0.986, p<0.001<br>[0.985, 0.987] | Excellent |

**Fig 7.**
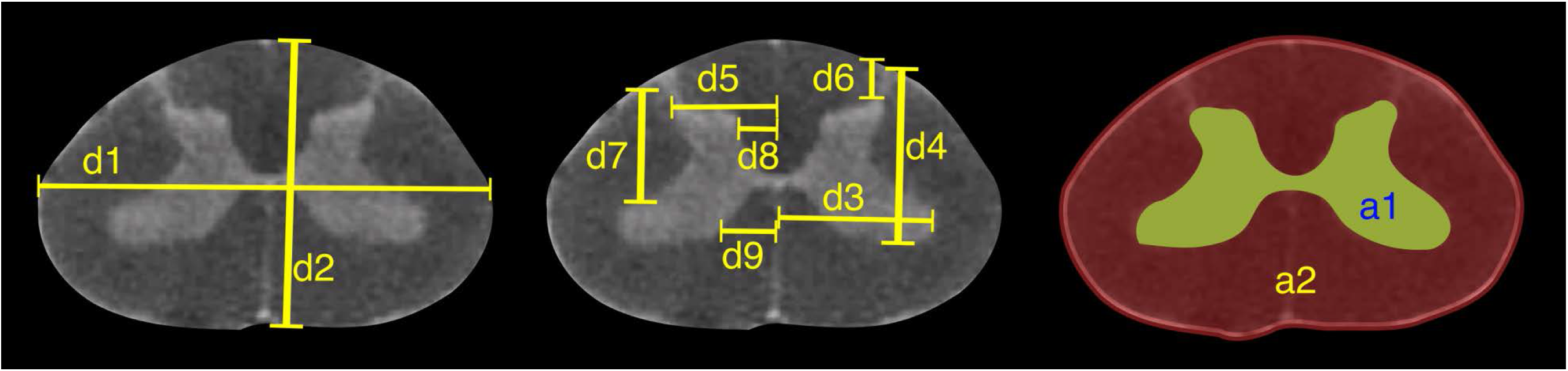
MRI-based morphometric measurements of the lumbosacral spinal cords of rat, cat, pig, monkey, and human. d1 and d2 represent the width and height of the spinal cord, respectively. d3 and d9 are the mediolateral distances from the midline to the lateral and medial boundaries of the ventral horn. Similarly, d5 and d8 are the mediolateral distances from the midline to the lateral and medial boundaries of the dorsal horn. d7 and d4 are the depth of the dorsal and ventral boundaries of the ventral horn from the spinal cord surface. d6 is the depth of the dorsal boundary of the dorsal horn. a2 and a1 represent the cross-sectional areas of the spinal cord and the grey matter, respectively.

**Fig. 8.**
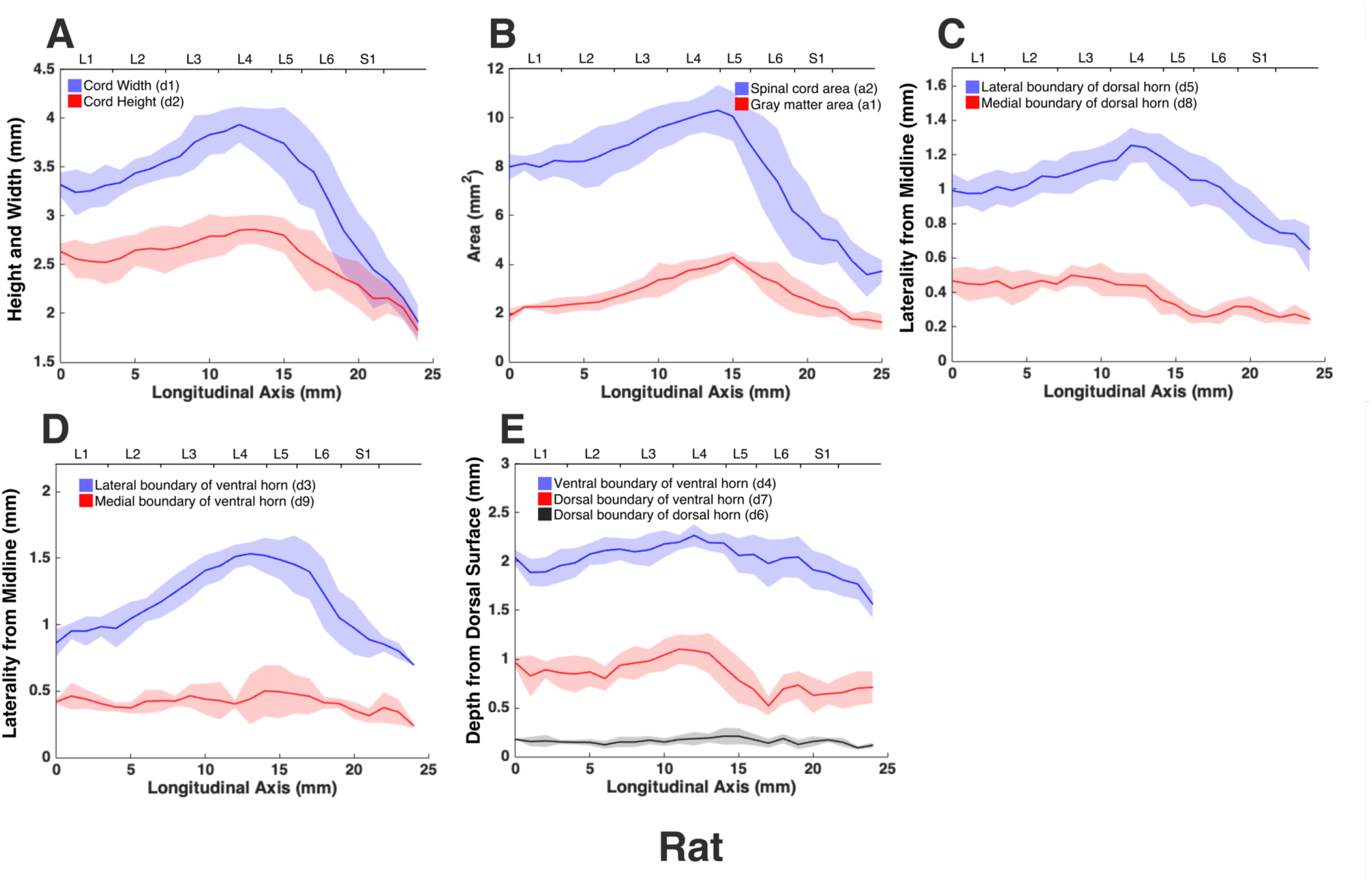
Measurements of cross-sectional dimensions of the spinal cord and locations of the ventral and dorsal horns of rat spinal cord. Presented data are based on n=6 specimens. Solid lines represent the mean and the shaded regions represent the mean ± standard deviation. A) Spinal cord width and height (d1 and d2, respectively in Fig. 7, left) across the lumbosacral cord. B) Cross-sectional areas of the spinal cord and grey matter (a1, a2 in Fig. 7, right) across the lumbosacral cord. C) Lateral distances from midline of dorsal horn boundaries (d5, d8 in Fig. 7, middle). D) Lateral distances from midline of ventral horn boundaries (d3, d9 in Fig. 7, middle). E) Depth from the dorsal surface of the spinal cord of the ventral and dorsal horn boundaries (d4, d7, d6, in Fig. 7, middle).

**Fig. 9.**
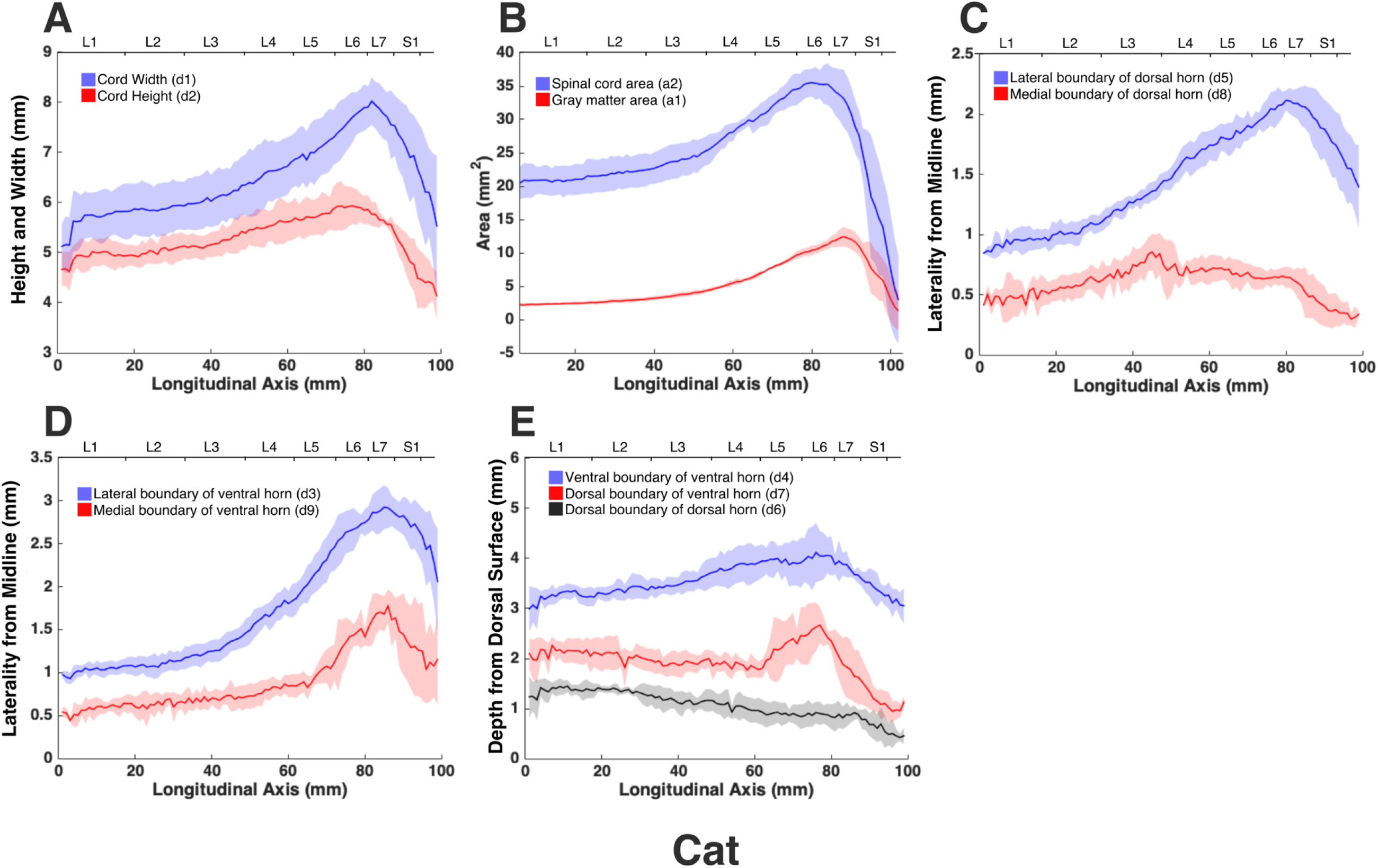
Measurements of cross-sectional dimensions of the spinal cord and locations of the ventral and dorsal horns of cat spinal cord. Presented data are based on n=6 specimens. Solid lines represent the mean and the shaded regions represent the mean ± standard deviation. A) Spinal cord width and height (d1 and d2, respectively in Fig. 7, left) across the lumbosacral cord. B) Cross-sectional areas of the spinal cord and grey matter (a1, a2 in Fig. 7, right) across the lumbosacral cord. C) Mediolateral distances from midline to the dorsal horn boundaries (d5, d8 in Fig. 7, middle). D) Mediolateral distances from midline to the ventral horn boundaries (d3, d9 in Fig. 7, middle). E) Depth from the dorsal surface of the spinal cord of the ventral and dorsal horn boundaries (d4, d7, d6, in Fig. 7, middle).

**Fig. 10.**
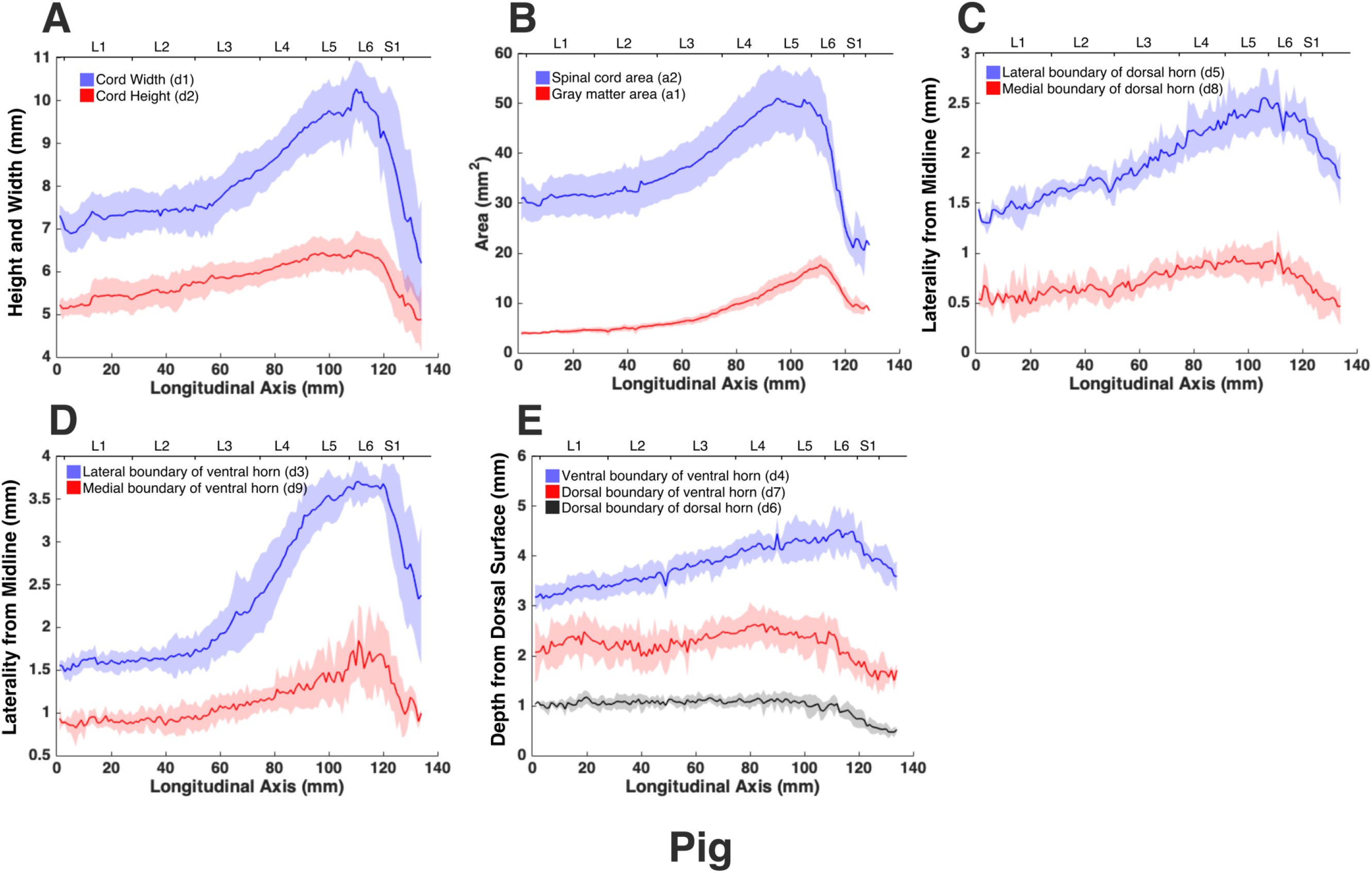
Measurements of cross-sectional dimensions of the spinal cord and locations of the ventral and dorsal horns of pig spinal cord. Presented data are based on n=6 specimens. Solid lines represent the mean and the shaded regions represent the mean ± standard deviation. A) Spinal cord width and height (d1 and d2, respectively in Fig. 7, left) across the lumbosacral cord. B) Cross-sectional areas of the spinal cord and grey matter (a1, a2 in Fig. 7, right) across the lumbosacral cord. C) Mediolateral distances from midline to the dorsal horn boundaries (d5, d8 in Fig. 7, middle). D) Mediolateral distances from midline to the ventral horn boundaries (d3, d9 in Fig. 7, middle). E) Depth from the dorsal surface of the spinal cord of the ventral and dorsal horn boundaries (d4, d7, d6, in Fig. 7, middle).

**Fig. 11.**
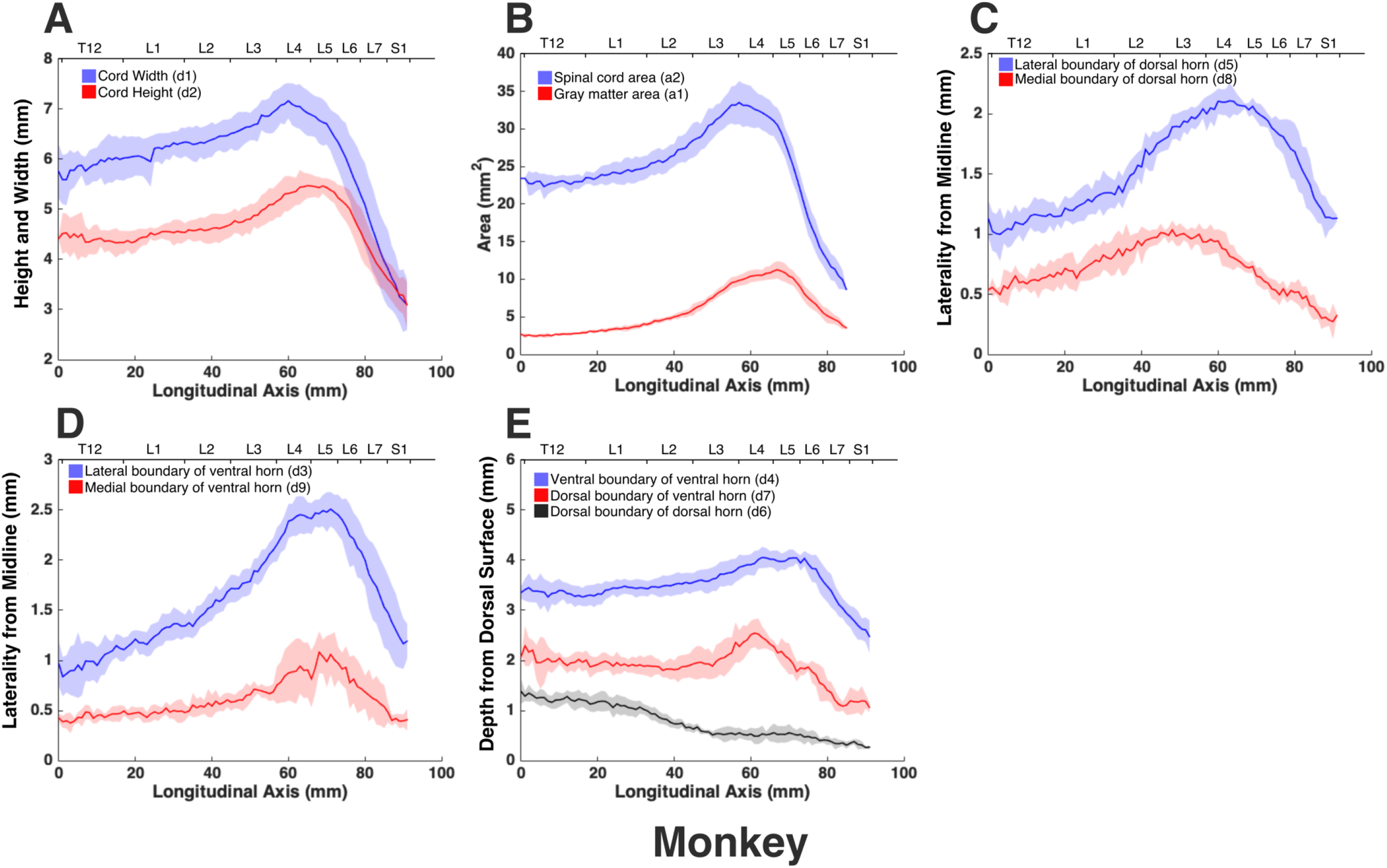
Measurements of cross-sectional dimensions of the spinal cord and locations of the ventral and dorsal horns of monkey spinal cord. Presented data are based on n=6 specimens. Solid lines represent the mean and the shaded regions represent the mean ± standard deviation. A) Spinal cord width and height (d1 and d2, respectively in Fig. 7, left) across the lumbosacral cord. B) Cross-sectional areas of the spinal cord and grey matter (a1, a2 in Fig. 7, right) across the lumbosacral cord. C) Mediolateral distances from midline to the dorsal horn boundaries (d5, d8 in Fig. 7, middle). D) Mediolateral distances from midline to the ventral horn boundaries (d3, d9 in Fig. 7, middle). E) Depth from the dorsal surface of the spinal cord of the ventral and dorsal horn boundaries (d4, d7, d6, in Fig. 7, middle).

**Fig. 12.**
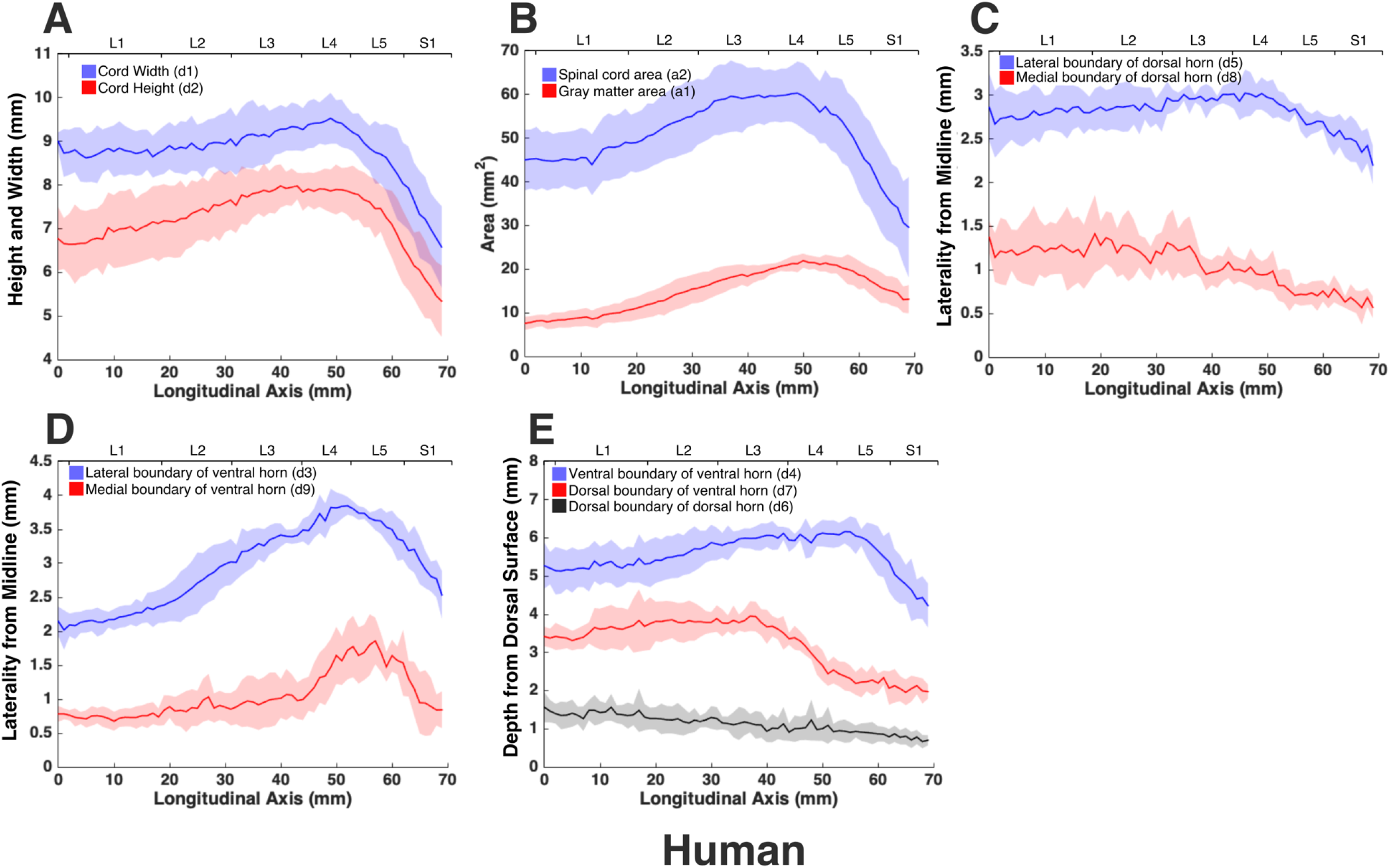
Measurements of cross-sectional dimensions of the spinal cord and locations of the ventral and dorsal horns of human spinal cord. Presented data are based on n=6 specimens. Solid lines represent the mean and the shaded regions represent the mean ± standard deviation. A) Spinal cord width and height (d1 and d2, respectively in Fig. 7, left) across the lumbosacral cord. B) Cross-sectional areas of the spinal cord and grey matter (a1, a2 in Fig. 7, right) across the lumbosacral cord. C) Mediolateral distances from midline to the dorsal horn boundaries (d5, d8 in Fig. 7, middle). D) Mediolateral distances from midline to the ventral horn boundaries (d3, d9 in Fig. 7, middle). E) Depth from the dorsal surface of the spinal cord of the ventral and dorsal horn boundaries (d4, d7, d6, in Fig. 7, middle).

In all species, moving from the rostral to the caudal end of the lumbosacral cord, the size of the spinal cord and the gray matter increase until they reach a peak, after which they decrease. The point at which the cord’s width (d1 in Fig. 7) reaches its peak value, is further referred to as ‘peak cord size’ or ‘PCS’ located in the enlargement. At the PCS, the spinal cord is 4.0 ± 0.2 mm wide (dimension d1 in Fig. 7, left) and 2.94 ± 0.2 mm high (dimension d2) in rats, 8.0±0.5 mm wide x 6.0±0.4 mm high in cats, 10.1±0.6 mm wide x 6.6 ± 0.3 mm high in pigs, 7.2±0.4 mm wide x 5.6±0.2 mm high in monkeys, and 9.6±0.6 mm wide x 8.2±0.5 mm high in humans (Fig. 13). In all species, the width of the spinal cord is larger than its height everywhere in the lumbosacral cord (Fig. 8–12). The cord’s aspect ratio (width/height or d1/d2 at PCS) is largest in pigs (1.54±0.06), followed by rats and cats (1.35±0.03 and 1.340±0.13, respectively), monkeys (1.29±0.09), and humans (1.18±0.06).

**Fig. 13.**
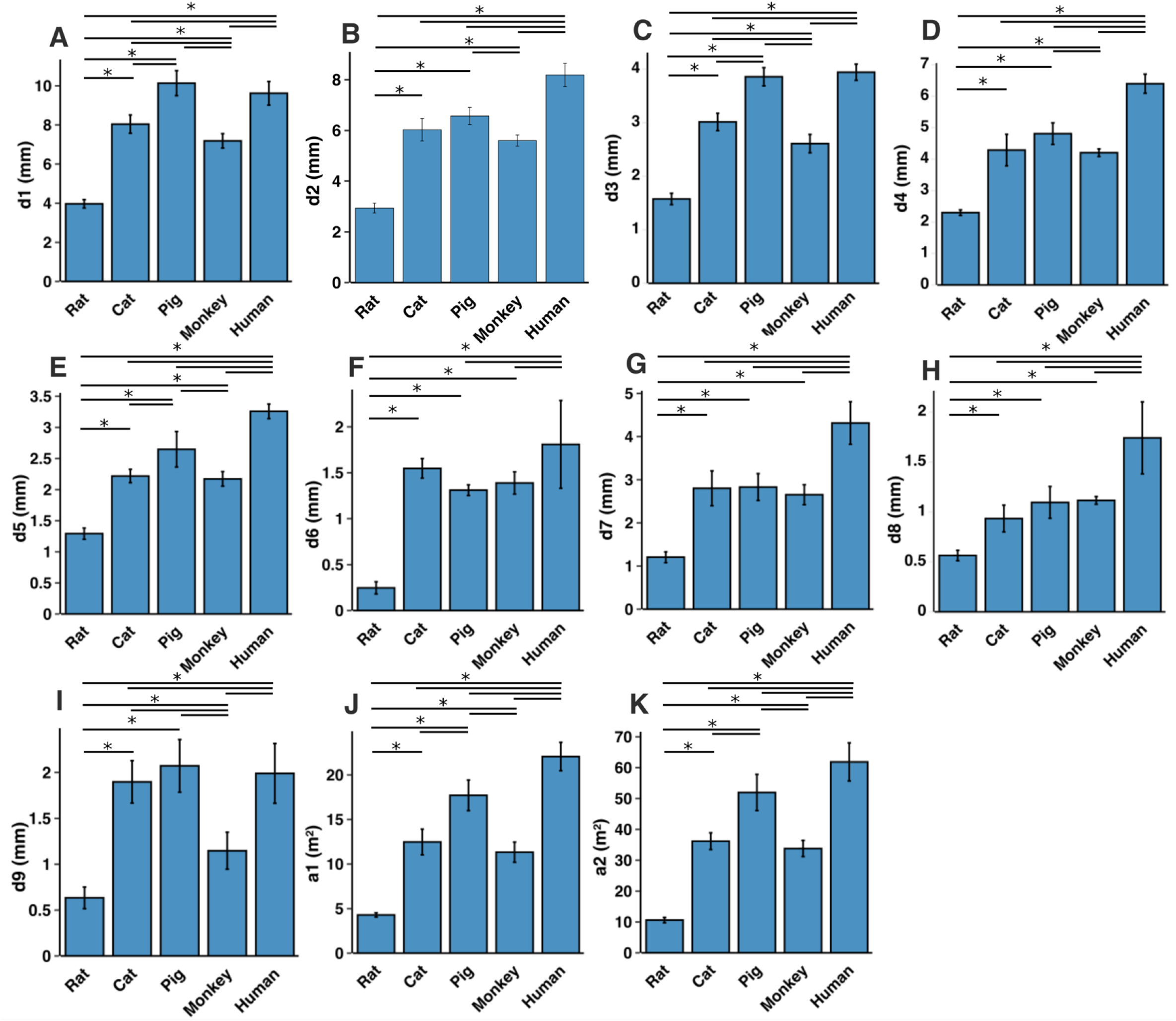
Comparison of the spinal cord dimensions and the locations of the dorsal and ventral horns in the spinal cords of rats, cats, pigs, monkeys, and humans. Graphs show the comparison of the values for each parameter at the Peak Cord Size (PCS) across species. Bars represent the mean and the error bars show the standard deviation of the mean. Parameters are those shown in Fig. 7 A) d1. B) d2. C) d3. D) d4. E) d5. F) d6. G) d7. H) d8. I) d9. J) a1. K) a2.

A comparison between the areas of the spinal cords across species shows that humans have the largest cord with a peak total cross-sectional area (a2 in Fig. 7, right) of 61.9±6.2 mm^2^ (Fig. 13). The size of the pigs’ spinal cord is smaller than that of the humans (p=0.003) and larger than that of the other species (p<0.001 for all comparisons) with a peak total area of 52.0±5.9 mm^2^. Monkey and cat cords are similar in size (p=0.863) with peak total area of 33.8±2.6 mm^2^ and 36.2±2.7 mm^2^, respectively. Rats have substantially smaller cords (p<0.001 for all comparisons) with peak total area of 10.6±0.9 mm^2^. A similar trend also exists for the peak total cross-sectional area of the grey matter (a1 in Fig. 7, right). The peak total cross-sectional area of the gray matter was 22.1±1.6 mm^2^, 17.70±1.7 mm^2^, 12.48±1.4 mm^2^, 11.33±1.1 mm^2^, and 4.29±0.2 mm^2^ for humans, pigs, cats, monkeys, and rats, respectively (Fig. 13). Moving caudally from the rostral end of the enlargement, the ratio of the total area of gray matter to that of the white matter (a1/a2) consistently increases in all species (Fig. S6).

At the PCS, the dorsal horns are on average 0.25±0.07 mm deep (d6 in Fig. 7, middle) relative to the dorsal surface of the cord, and 1.29±0.09 mm lateral to the midline (d5) in rats, 1.54±0.11 mm deep and 2.22±0.11 mm lateral to the midline in cats, 1.31±0.06 mm deep and 2.65±0.29 mm lateral to the midline in pigs, 1.39±0.12 mm deep and 2.17±0.12 mm lateral to the midline in monkeys, and 1.81±0.48 mm deep and 3.26±0.12 mm lateral to the midline in humans. At PCS, the ventral horns are on average 2.30±0.09 mm deep (d4) and 1.57±0.11 mm lateral to the midline (d3) in rats, 4.29±0.50 mm deep and 3.01±0.16 mm lateral to the midline in cats, 4.81±0.34 mm deep and 3.86±0.17 mm lateral to the midline in pigs, 4.20±0.12 mm deep and 2.61±0.17 mm lateral to the midline in monkeys, and 6.40±0.30 mm deep and 3.94±0.15 mm lateral to the midline in humans.

In rats, the dorsal horns are shallower relative to the cord height (d6/d2 in Fig. 7) compared to other species. The shallowest depth of the dorsal horns on average ranges between 3-10%, 7-31%, 9-23%, 5-32%, and 8-26% of the cord height in rats, cats, pigs, monkeys and humans, respectively. In rats, monkeys and humans, the ventral horns extend deeper in the spinal cord relative to their cord height than the other species. The largest depth of the ventral horns on average ranges between 72-90%, 61-79%, 57-82%, 70-84%, and 69-87% of the cord height (d4/d2) in rats, cats, pigs, monkeys and humans, respectively. In rats, monkeys and humans, the dorsal horns also extend more laterally relative to their cord width, than the other species. The largest laterality of each of the dorsal horns on average ranges between 28-35%, 15-30%, 18-32%, 17-38%, and 28-40% of the width of the spinal cord (d5/d1) in rats, cats, pigs, monkeys and humans, respectively.

The laterality of the ventral horns relative to their cord width (d3/d1 in Fig. 7) is similar across species. The largest laterality of each of the ventral horns on average ranges between 27-42% (peak located 4.2±1.5 mm caudal to PCS), 17-43% (peak located 12.3±6.9 mm caudal to PCS), 20-43%(peak located 12.7±10.6 mm caudal to PCS), 15-43% (peak located 22.5±3.3 mm caudal to PCS), and 23-44% (peak located 11.5±2.7 mm caudal to PCS) of the width of the spinal cord in rats, cats, pigs, monkeys and humans, respectively. Values of all parameters at the PCS are compared across species in Fig. 13.

## Discussion

In this paper we presented a high-resolution comparative atlas of the spinal cord with detailed anatomical information about the lumbosacral region in five mammalian species. Results include comparative location, length, cross-sectional area, and morphology of the grey and white matter of intact spinal cords in rats, cats, pigs, monkeys, and humans. To the best of our knowledge, this is the first comprehensive comparative spinal cord atlas for animal species that are common subjects of basic and translational research. Not only can this atlas be used as a reference for the typical anatomy of the spinal cord in studies of spinal cord pathologies and injuries, but it also can guide researchers in selecting appropriate animal models in various stages of basic and translational research [38]. A comparison between the obtained species atlases reveals that the lumbosacral spinal cord of pigs is the most similar in size to that in humans. This includes the length and cross-sectional area of the spinal cord as well as the locations of the dorsal and ventral horns. Cats and monkey spinal cords are also similar in size and smaller than that of the humans. Rats have the smallest spinal cord.

To validate our findings, we compared the sizes of the human and pig spinal cords to those reported in the literature based on a combination of imaging and histological records. Frostell et al [39] reviewed and combined the findings of 11 studies of the segmental sizes of the human spinal cord obtained through histology and imaging. They estimated the width (d1) and height (d2) of spinal cord segments L1-S1 to range on average between 8.4-9.4 mm and 6.7-7.5 mm, respectively. These are consistent with our measurements of width, ranging on average between 6.6-9.5 mm, and height ranging on average between 5.3-7.9 mm for this spinal cord region. Cuellar et al [40], studied the spinal cord anatomy of pigs histologically, and reported spinal cord width (d1) across the L1-L6 segments. On average, cord width ranged between 7.5-9.4 mm in their study, which is consistent with our findings of cord width, ranging on average between 6.9-10.3 mm.

Knowledge about the dimensions of the spinal cord is important for investigations of treatments such as stem cell therapies [41] and spinal cord neuroprostheses [35], [42]. Knowing the dimensions of targeted regions of the spinal cord is essential for surgical planning and successful delivery of cells, drugs, electrodes or optrodes to their intraparenchymal targets. This information can guide the technological design of the necessary implants [33], [42], [43] and delivery apparatuses [22], [44], [45]. Geometrical dimensions are also important for informed interpretation of cell migration and drug perfusion in preclinical models and their implications for human translation.

As an example, intraspinal microstimulation (ISMS) [17], [18], [33] is a spinal cord neuroprosthesis for restoring mobility after spinal cord injury that involves the implantation of fine microelectrodes into the ventral horns of the spinal cord to activate functional spinal motor networks. ISMS microelectrodes are typically implanted perpendicularly to the major axis of the cord along the anteroposterior (dorsoventral) axis [22], [42], and target regions within Rexed lamina IX of the spinal cord [35], [46], [47]. Therefore, technological design and surgical implantation of ISMS implants requires species- and segment-specific knowledge about the location and dimensions of the ventral horns. Created atlases and the presented morphological measurements address this need, by informing the location and dimensions of the targeted ventral horns in various species (d3, d4, d7, and d9, shown in Fig. 7).

The comparative atlas created in this study was based on images of perfused spinal cord specimen which may have experienced some shrinkage due to the perfusion process. Siefert et al [48] quantified the level of shrinkage in human spinal cords 1 day and 31 days post formalin fixation and found that the cross-sectional size of the spinal cords shrinks by an average of 4%, and 4.4%, respectively. Therefore, an appropriate correction factor should be taken into consideration when absolute measurement values are needed. It is also important to consider the correlation of age and cord dimensions when using absolute measurement values. Papinutto et al [49] reported that the cross-sectional area of the human spinal cord decreases with aging. The average age of the donors of our study was 83±8 years (table 1).

The focus of this work was on creating high-resolution atlases for the lumbosacral spinal cord in species commonly used in basic and translational research. Future investigations should consider studying other regions of the spinal cord, eventually resulting in complete anatomical documentation for the entire spinal cord. Similar methodology can also be applied to develop spinal cord atlases for various pathologies to enhance our understanding of these conditions.

## Materials and Methods

All procedures were performed according to protocols approved by the animal care and use committees at the Universities of Alberta and Washington, and the human research ethics board at the University of Alberta.

### Spinal cord extraction

Rats, cats, and monkeys, were perfused transcardially with 4% formaldehyde solution to achieve tissue fixation. In pigs, spinal cord specimens were extracted from freshly euthanized animals and immediately placed in 4% formaldehyde solution for fixation. Human cadavers had been embalmed by the Division of Anatomy at the University of Alberta, prior to tissue extraction. In order to extract spinal cord specimens, multi-level laminectomies were performed to expose spinal cord segments and spinal nerve roots from T12-S3. The spinal nerve roots were identified and marked. Spinal cord specimens were removed with the spinal nerve roots attached and stored in 4% formaldehyde solution.

### Measurements of spinal cord segments

Prior to MR imaging, boundaries of the spinal cord segments were identified and marked with glass tubes (3-5 mm in length, 170 μm in diameter, Wale Apparatus Company, Hellertown, USA). In this procedure, the dura mater and arachnoid were opened, and spinal cord segments were identified based on the location of the dorsal rootlets entering the spinal cord (dorsal root entry zones (DREZ)) (Fig. S7). The most rostral and most caudal rootlets entering the cord for each segment were carefully tracked back to their respective roots using a dissection microscope. Boundaries of spinal segments were defined as the middle point between neighboring DREZs. For each identified boundary, a glass tube was inserted in the transverse plane of the spinal cord parenchyma. The glass tubes were visible in MRIs. The rostrocaudal lengths of all identified spinal cord segments were measured using a ruler.

### Imaging

The spinal roots were carefully cut from the cords prior to imaging. Spinal cord specimens were then placed in glass tubes filled with Fluorinert (FC-770, Milipore Sigma, Darmstadt, Germany) for imaging. MR images of specimen from rats, cats, pigs, and humans were acquired on a 4.7T Varian MRI scanner (Varian Inc., Palo Alto, USA) in the Peter S Allen MR Research Center at the University of Alberta. A 3D gradient echo sequence was used (TR/TE = 39.7 ms/28 ms, 1 average, FOV= 40 mm x 40 mm x 60 mm, 60 transverse slices, 1 mm slice thickness) with a resolution of 0.125×0.125×1mm. These scans were conducted using a 38mm diameter, 35 mm long volume coil with the Litz design (Doty Scientific, Columbia, USA). In order to acquire images of the entire spinal cord specimen that were longer than this coil, a custom-made setup was used to translate the specimen horizontally within the coil, by precise steps.

MR images of the monkey spinal cords were acquired on a 3T Philips Achieva MRI scanner (Best, Netherlands) at the University of Washington Diagnostic Imaging Sciences Center. All scans used an eight channel Philips wrist coil. Two scans were acquired. First, a merged (four echo) fast field echo (mFFE) sequence, with TR/TE/deltaTE = 126ms/6.3ms/11.7ms, 1 average. Secondly, a T1-weighted MPRAGE scan with TR/TE/TFE factor = 21ms/5.8ms/80, TI = 950ms, a shot interval = 2500ms, and four averages. Both scans had a FOV of 64 mm x 48 mm x 112 mm, 112 transverse slices, and 1 mm slice thickness) with a resolution of 0.15×0.15×1mm.

### 3D Model

3D models of the lumbosacral spinal cords were generated based on the acquired MRIs. Transverse MR images of the spinal cords (one typical spinal cord per species) were traced and their coordinates were extracted, using a custom written program in Python (version 3.7, Python software foundation, Wilmington, DE, USA). Extracted coordinates were used to reconstruct 3D models.

### MRI-based measurements

The morphology of all spinal cords was quantified through measurements of distance and area as shown in Fig. 7. These parameters captured the size of the grey and white matter as well as the location of the dorsal and ventral horns in the spinal cords of the studied species. Each parameter for each specimen was measured manually by two raters (a total of 5 raters in the study) using ImageJ software (National Institute of Health). Raters were blinded to the scale of the image pixels as well as the species of the image sets. Measurements were then transferred to Matlab (version 2015a, MathWorks, Natick, MA, USA) for group analysis and data visualization. In order to average the measured parameters for each species (Fig. 8–12), across 6 specimens over their entire length, curves were aligned so that their PCS locations (where d1 was maximum) lines up with x=0 (Fig. S1–S5).

### Statistical analysis

Comparisons were made between spinal cord measurements across spinal segments and species using one-way ANOVAs and Tukey honest significant difference post-hoc tests. In order to assess the inter-rater reliability of the MRI-based measurements, ICC coefficients were calculated [50]. ICC calculations were done for each species based on the one-way random model and ICC-type representing the mean of 2 raters [51]. Values between 0.5 and 0.75, 0.75 and 0.9, larger than 0.9, were assessed as moderate, good, and excellent reliability, respectively [51]. All statistical analyses were performed using RStudio software (version 1.2.5019, RStudio Inc., Boston, USA). For each species, all measurements are reported as mean ± standard deviation.

## Acknowledgments

The authors would like to acknowledge the cadaveric donors for their valuable contribution to this research. We thank Jason Papirny for facilitating access to human cadavers through the Division of Anatomy at the University of Alberta. We also thank the staff of the Ray Rajotte Surgical Medical Research Institute at the University of Alberta for facilitating access to swine tissue. Thanks to the Washington National Primate Research Centre staff for facilitating access to spinal cord tissue from non-human primates. We also thank Chris English, Rob Robinson, Bethany Kondiles, and Rod Gramlich for assisting with the extraction, perfusion and dissection of the non-human primate tissue, and for fabricating an MR compatible setup that allowed horizontal translation of the imaged samples inside the magnet.

This work was funded in part by the Canadian Institutes of Health Research, the Canada Foundation for Innovation and the National Institute of Health (grant R01NS012542). AT was supported by a CIHR Vanier Canada Graduate Scholarship, an Alberta Innovates Health Solutions Studentship, and a Queen Elizabeth II Graduate Scholarship. VKM is a Canada Research Chair (Tier 1) in Functional Restoration.

## Supplementary Tables

**Table S1.** Segmental ranges and length of the lumbosacral enlargement in individual samples

| Sample ID | Species | Spinal Cord Levels of the Lumbosacral Enlargement | Length of the lumbosacral enlargement (mm) |
| --- | --- | --- | --- |
| R1 | Rat | L2-S1 | 17 |
| R2 | Rat | L3-L6 | 12 |
| R3 | Rat | L3-S1 | 12 |
| R4 | Rat | L3-L6 | 10 |
| R5 | Rat | L3-S1 | 12 |
| R6 | Rat | L3-S1 | 11 |
| C1 | Cat | L4-S1 | 35 |
| C2 | Cat | L4-S2 | 37 |
| C3 | Cat | L4-S1 | 33 |
| C4 | Cat | L4-S1 | 33 |
| C5 | Cat | L4-S1 | 34 |
| C6 | Cat | L4-S2 | 34 |
| P1 | Pig | L3-S2 | 69 |
| P2 | Pig | L3-S1 | 76 |
| P3 | Pig | L3-S2 | 67 |
| P4 | Pig | L3-S1 | 60 |
| P5 | Pig | L3-S1 | 62 |
| P6 | Pig | L3-S1 | 66 |
| M1 | Monkey | L2-L7 | 31 |
| M2 | Monkey | L3-S1 | 32 |
| M3 | Monkey | L3-L7 | 30 |
| M4 | Monkey | L3-S1 | 38 |
| M5 | Monkey | L2-L7 | 33 |
| M6 | Monkey | L2-S2 | 45 |
| H1 | Human | T12-S1 | 50 |
| H2 | Human | T12-S2 | 64 |
| H3 | Human | T12-S2 | 56 |
| H4 | Human | L1-S1 | 57 |
| H5 | Human | L1-S1 | 56 |
| H6 | Human | L2-S2 | 63 |

## Supplementary Figures

**Fig. S1.**
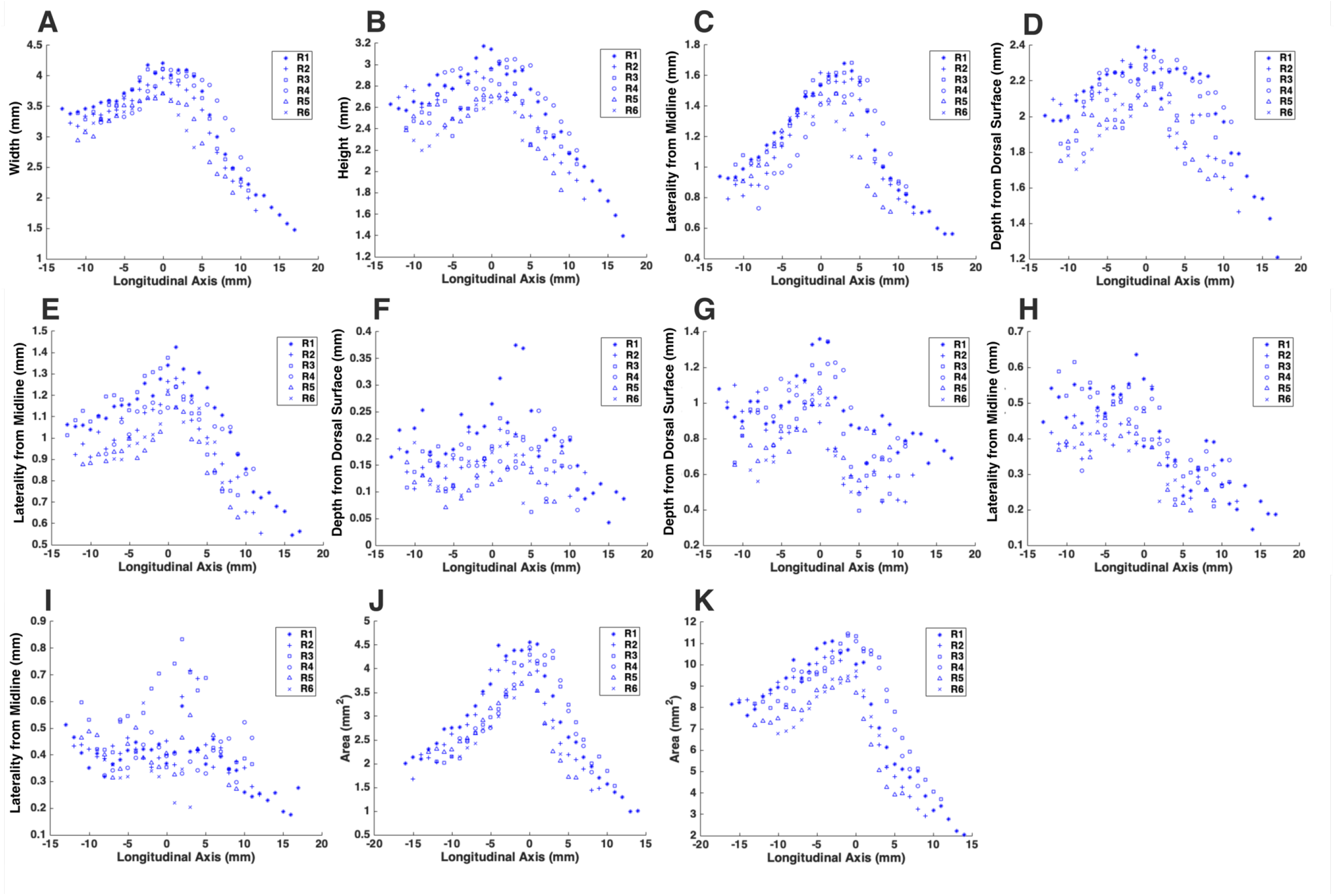
Cross-sectional dimensions of the spinal cord and locations of the ventral and dorsal horns of rat cords. Presented data are based on n=6 specimens per species. Different symbols represent different specimens within a species. Measurements are of the parameters shown in Fig. 7 across the length of the lumbosacral cord. All curves have been translated across the x-axis such that the cord’s PCS (where d1 is maximum) is at the origin (x=0). A) d1. B) d2. C) d3. D) d4. E) d5. F) d6. G) d7. H) d8. I) d9. J) a1. K) a2.

**Fig. S2.**
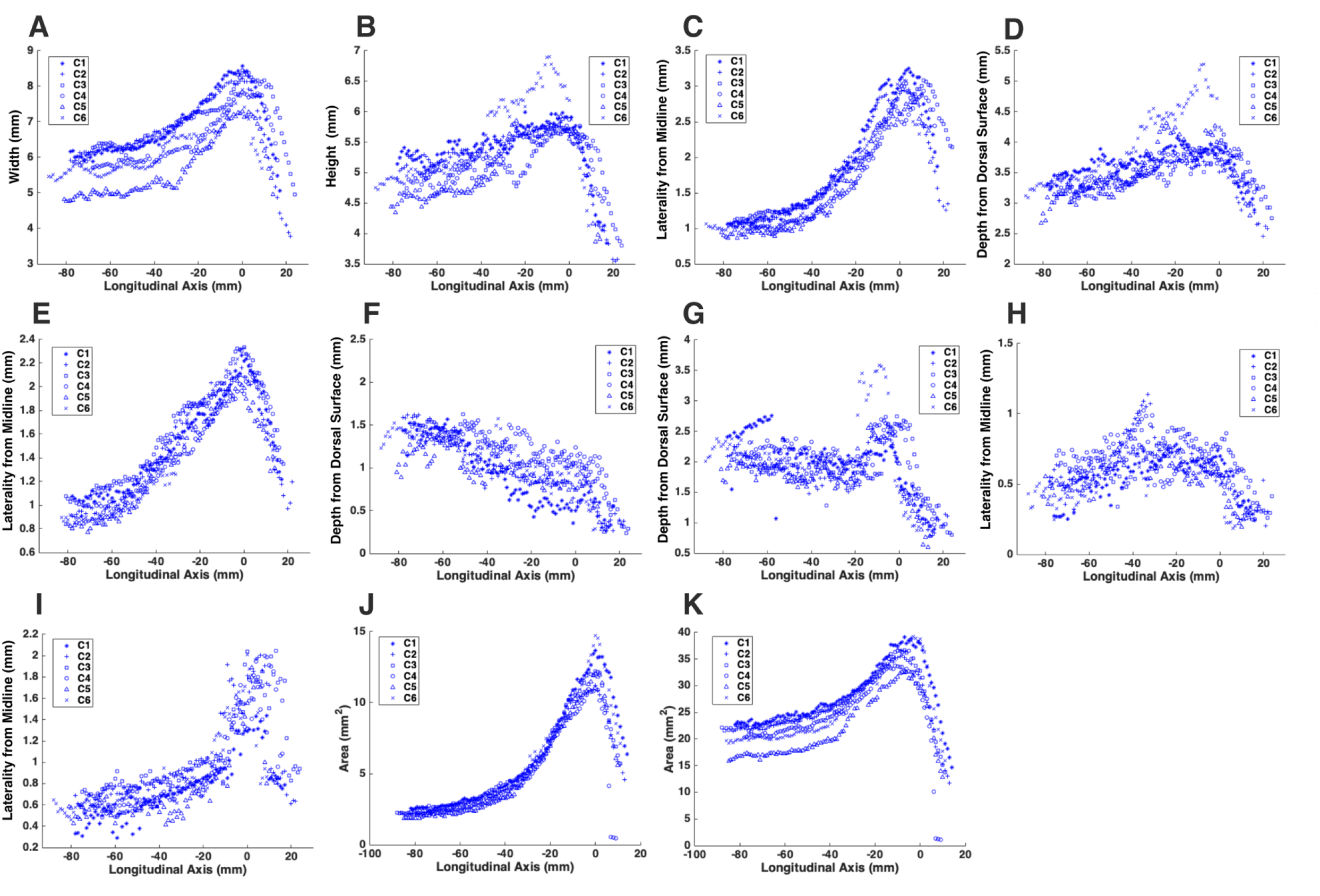
Cross-sectional dimensions of the spinal cord and locations of the ventral and dorsal horns of cat cords. Presented data are based on n=6 specimens per species. Different symbols represent different specimens within a species. Measurements are of the parameters shown in Fig. 7 across the length of the lumbosacral cord. All curves have been translated across the x-axis such that the cord’s PCS (where d1 is maximum) is at the origin (x=0). A) d1. B) d2. C) d3. D) d4. E) d5. F) d6. G) d7. H) d8. I) d9. J) a1. K) a2.

**Fig. S3.**
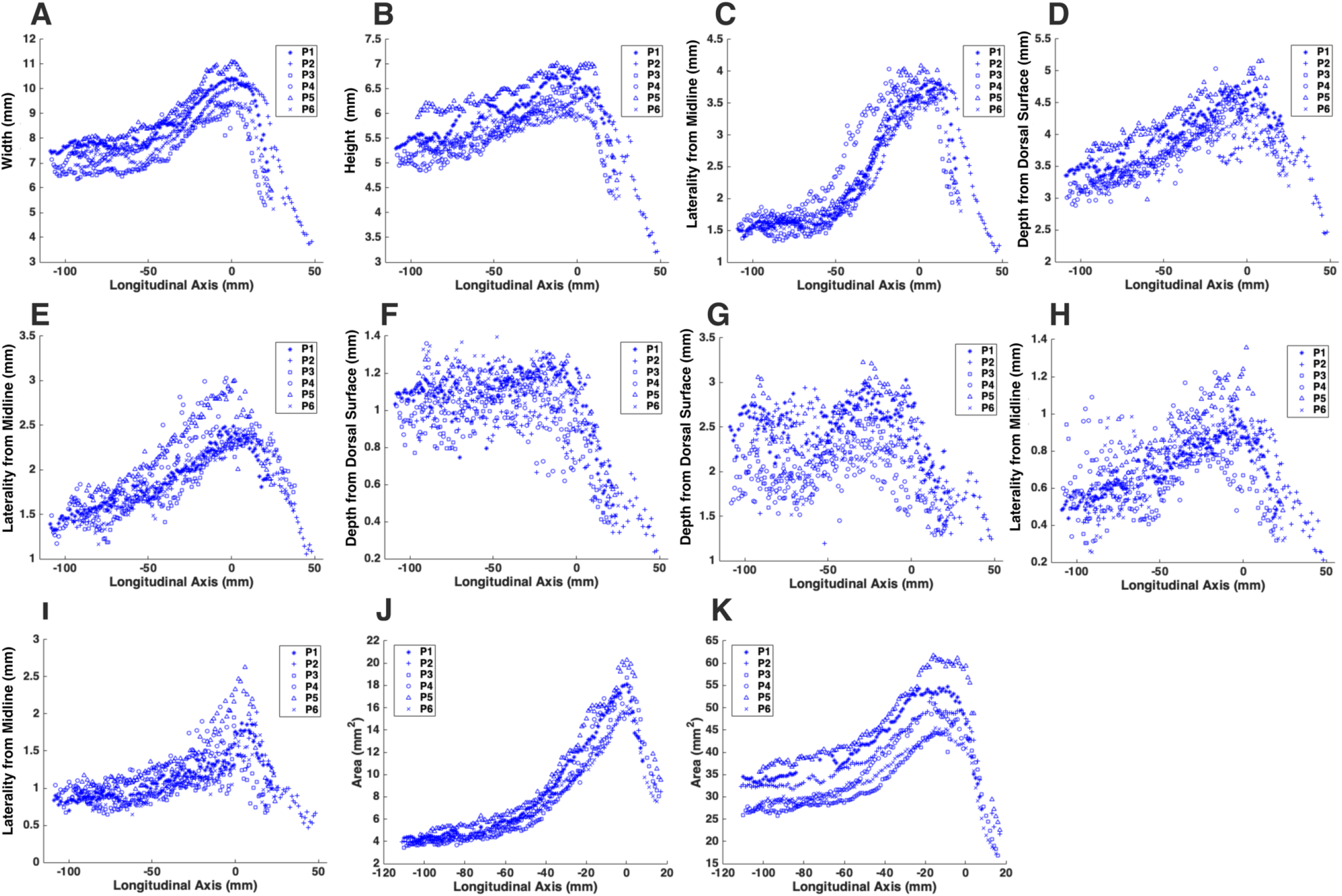
Cross-sectional dimensions of the spinal cord and locations of the ventral and dorsal horns of pig cords. Presented data are based on n=6 specimens per species. Different symbols represent different specimens within a species. Measurements are of the parameters shown in Fig. 7 across the length of the lumbosacral cord. All curves have been translated across the x-axis such that the cord’s PCS (where d1 is maximum) is at the origin (x=0). A) d1. B) d2. C) d3. D) d4. E) d5. F) d6. G) d7. H) d8. I) d9. J) a1. K) a2.

**Fig. S4.**
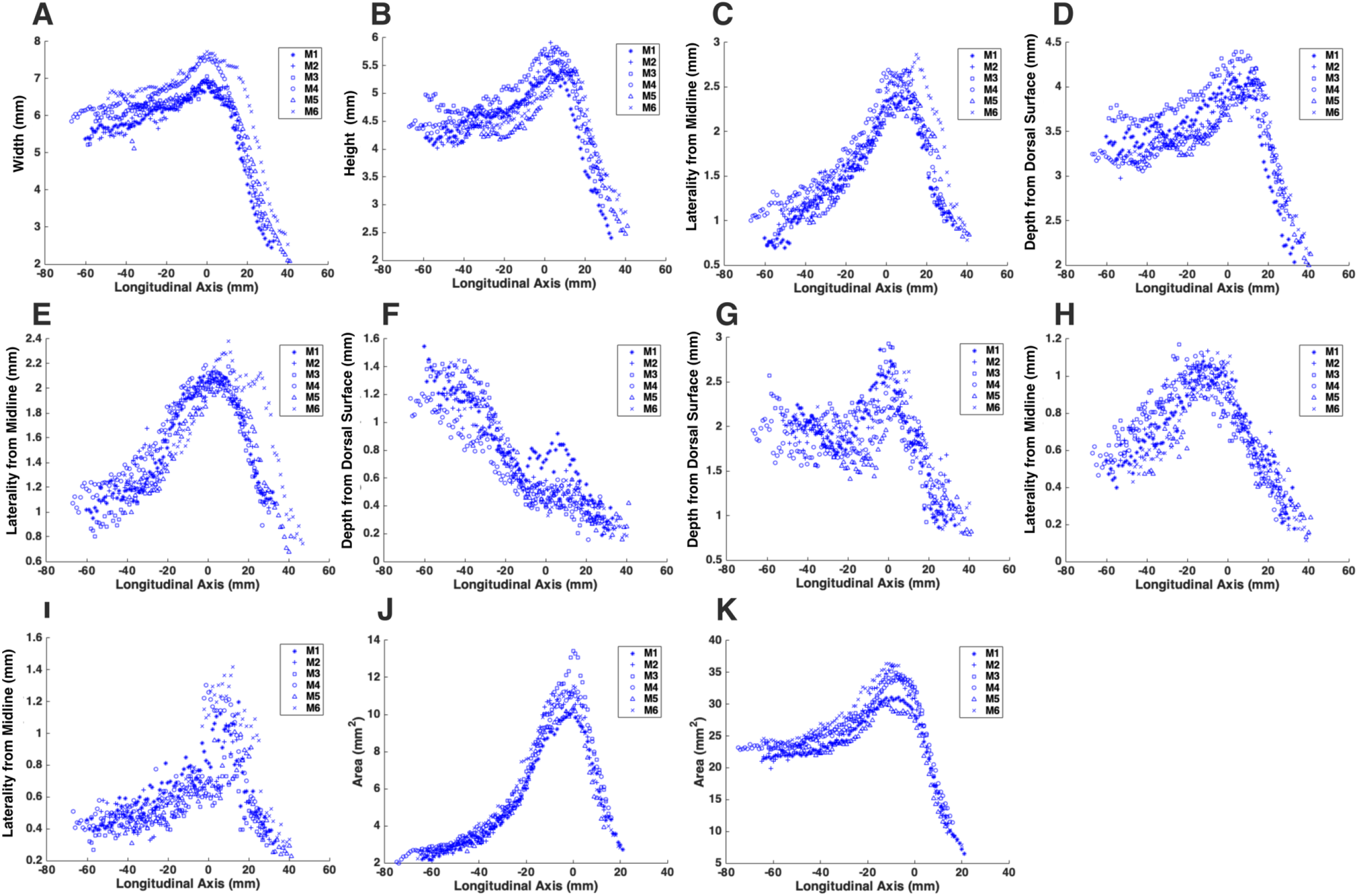
Cross-sectional dimensions of the spinal cord and locations of the ventral and dorsal horns of monkey cords. Presented data are based on n=6 specimens per species. Different symbols represent different specimens within a species. Measurements are of the parameters shown in Fig. 7 across the length of the lumbosacral cord. All curves have been translated across the x-axis such that the cord’s PCS (where d1 is maximum) is at the origin (x=0). A) d1. B) d2. C) d3. D) d4. E) d5. F) d6. G) d7. H) d8. I) d9. J) a1. K) a2.

**Fig. S5.**
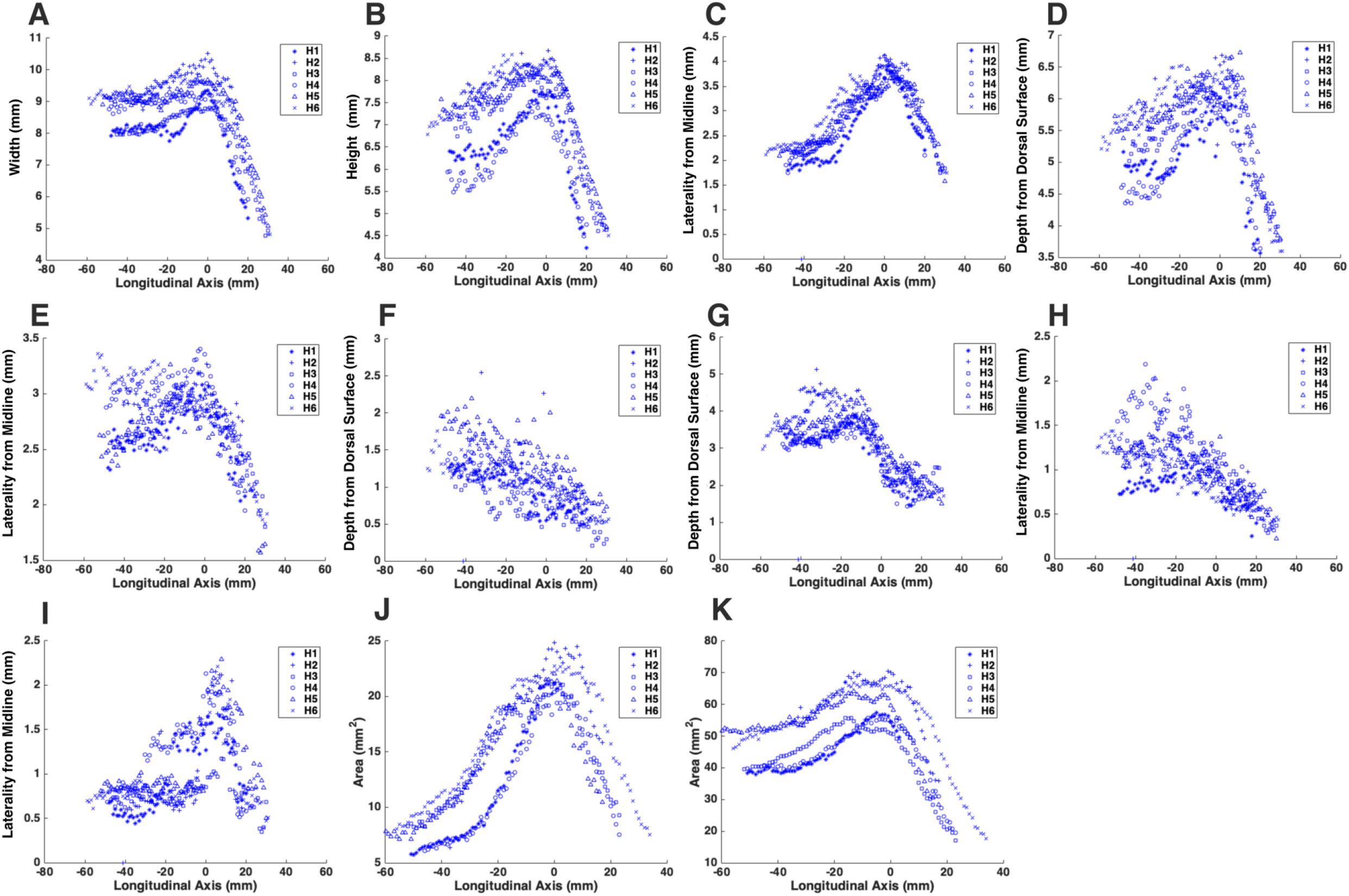
Cross-sectional dimensions of the spinal cord and locations of the ventral and dorsal horns of human cords. Presented data are based on n=6 specimens per species. Different symbols represent different specimens within a species. Measurements are of the parameters shown in Fig. 7 across the length of the lumbosacral cord. All curves have been translated across the x-axis such that the cord’s PCS (where d1 is maximum) is at the origin (x=0). A) d1. B) d2. C) d3. D) d4. E) d5. F) d6. G) d7. H) d8. I) d9. J) a1. K) a2.

**Fig. S6.**
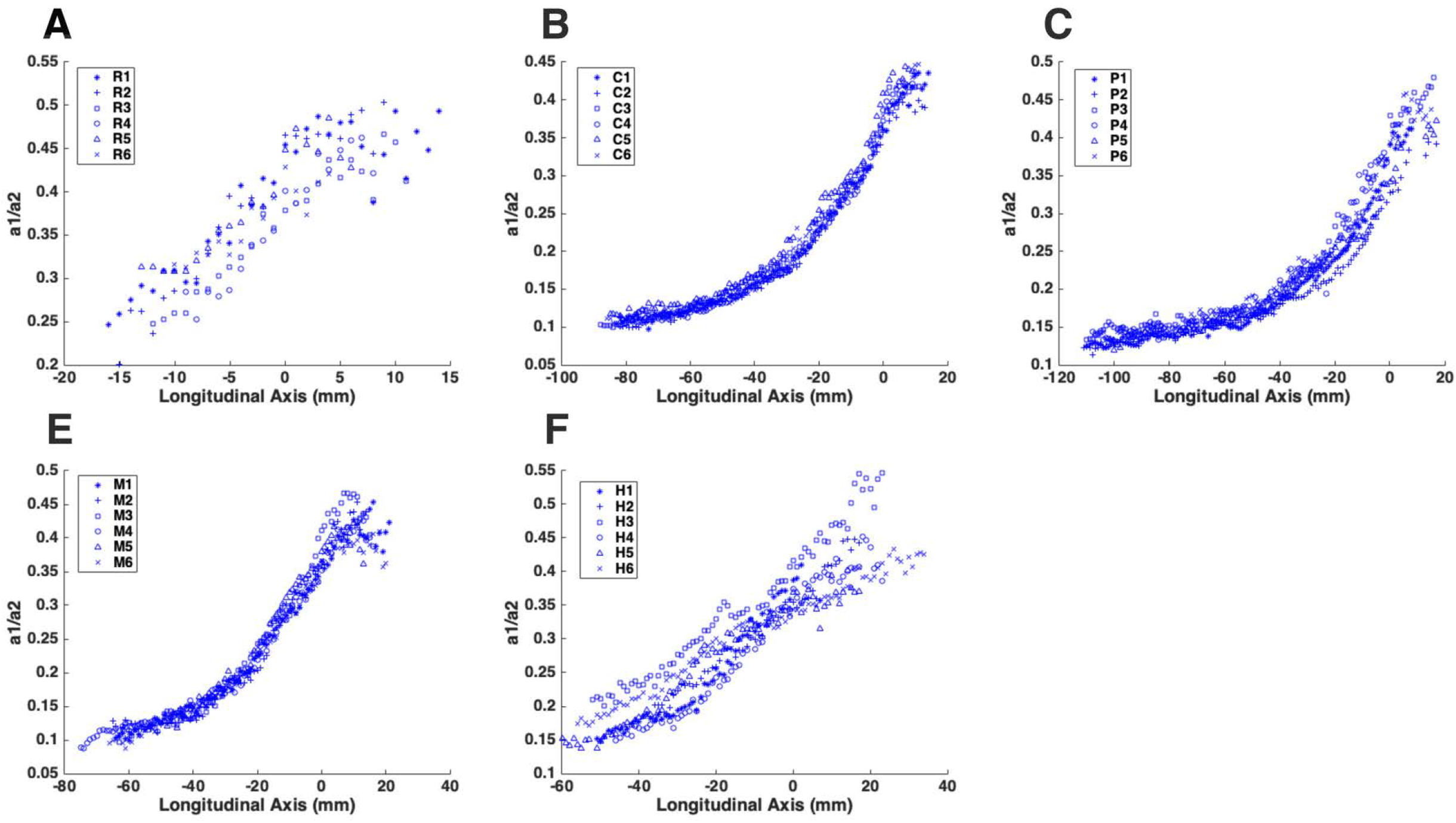
Ratio of the total area of the gray matter to that of the white matter (a1/a2) in all spinal cords of A) Rat, B) Cat, C) Pig, D) Monkey, E) Human. Presented data are based on n=6 extracted specimens per species. Different symbols represent different specimens. All curves have been translated across the x-axis so that each cord’s PCS (where d1 is maximum) is at the origin (x=0)

**Fig. S7.**
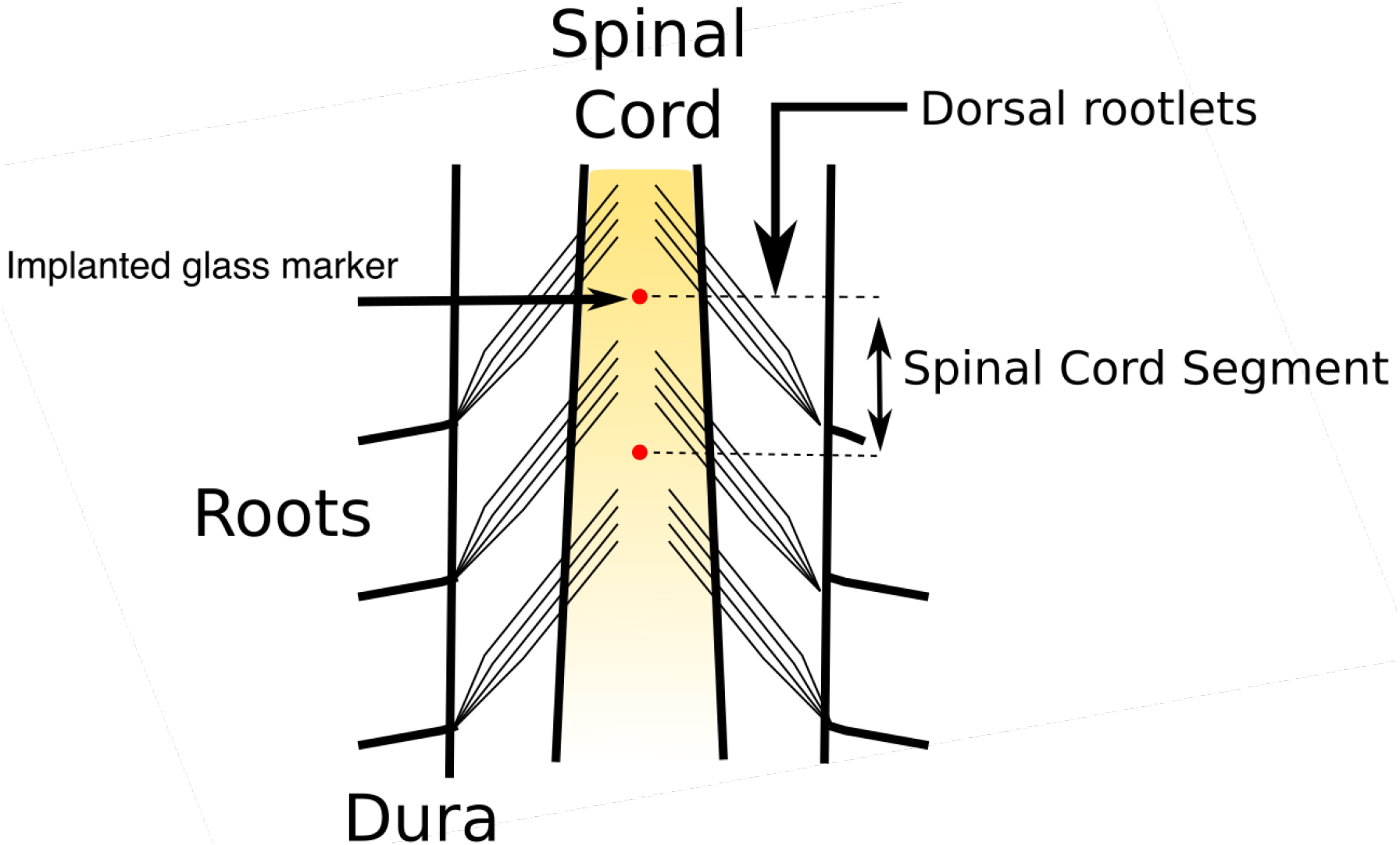
Spinal cord segment identification method (dorsal view). Spinal cord segments were identified based on the location of dorsal rootlets.

